# Deep phenotyping of ATDC5-derived in vitro cartilage organoids

**DOI:** 10.64898/2026.03.16.711783

**Authors:** Anna Klawonn, Stefan Tholen, Ilona Skatulla, Chiara M. Schröder, Sebastian J. Arnold, Oliver Schilling, Miriam Schmidts

**Affiliations:** Center for Pediatrics and Adolescent Medicine, University Hospital Freiburg, Freiburg, Germany; CIBSS - Centre for Integrative Biological Signalling Studies, University of Freiburg, Freiburg, Germany; Spemann Graduate School of Biology and Medicine (SGBM), Freiburg, Germany; Faculty of Biology, University of Freiburg, Freiburg, Germany; Institute for Surgical Pathology, Medical Center – University of Freiburg / Medical Faculty – University of Freiburg, Germany; Proteomics Platform – Core Facility (ProtCF), Medical Center – University of Freiburg / Medical Faculty – University of Freiburg, Germany; Institute of Experimental and Clinical Pharmacology and Toxicology, Faculty of Medicine, University of Freiburg, Germany

**Author notes:** corresponding author Dr. Miriam Schmidts, Department of Pediatrics, Adolescent Medicine and Neonatology Medical Center – University of Freiburg Faculty of Medicine, Breisacher Straße 86, 79106 Freiburg Germany.

**Keywords:** Chondrogenesis, Cartilage, ATDC5, Extracellular matrix, Proteomics

## Abstract

Cartilage is characterized by a highly specialized extracellular matrix (ECM) secreted by chondrocytes and limited self-regenerative capacity. In vivo investigations of chondrogenesis are limited by difficult and traumatic access, especially in humans. While it is known for decades that disturbances of chondrocyte differentiation and changed cartilage ECM composition cause severe skeletal phenotypes in vertebrates, a detailed molecular understanding of chondrogenesis and cartilage ECM formation is still missing, especially in the context of human genetic skeletal diseases.

ATDC5 cells, derived from AT805 mouse teratocarcinoma cells, have been used in the past to model chondrogenic differentiation, however, most studies have investigated few major cellular differentiation markers only so that the composition of the secreted ECM as well as effects on the ATDC5 transcriptome upon differentiation are still unclear. Here, we performed time-resolved transcriptomic and ECM proteomic analyses of differentiating ATDC5 cells. Both datasets confirmed the formation of a cartilage-like matrix with increasing expression of key chondrocyte genes over the course of differentiation. ECM proteomics further revealed a number of ECM components not previously reported in ATDC5 cells or the secreted ECM, encompassing collagens, proteoglycans, glycoproteins and other secreted factors. Overall, our findings provide a more detailed molecular characterization of ATDC5 chondrogenesis and highlight the potential of this model system for ECM-focused studies.

## 1. Introduction

Cartilage is a unique tissue characterized by a low cellular density and an abundant, highly specialized extracellular matrix (ECM) secreted by chondrocytes [1]. This matrix is predominantly composed of collagens and proteoglycans and is responsible for the characteristic mechanical properties of cartilage [2,3]. While collagen fibrils form a stable and tensile network [4], highly water-binding proteoglycans confer elasticity and resistance to compressive forces [5]. In addition, a wide range of essential non-collagenous proteins contribute to the interconnection of supramolecular structures, fibril assembly, matrix integrity, and cell-matrix interactions [6,7]. Understanding the process of chondrogenesis, ECM composition and homeostasis, and the interactions between chondrocytes and their surrounding matrix is of considerable interest. This knowledge is not only critical for elucidating defects in endochondral bone formation [8], but is also of importance for advancing human cartilage tissue engineering, as the treatment of cartilage injuries and degenerative diseases, such as osteoarthritis, remain a challenge in modern regenerative medicine [9].

Chondrogenesis is a multistep process that begins with the condensation of mesenchymal progenitor cells, followed by their differentiation into mature chondrocytes. During endochondral bone formation, these cells subsequently progress through a proliferating, then prehypertrophic stage and further differentiate into hypertrophic chondrocytes. Hypertrophic chondrocytes exit the cell cycle and contribute to the mineralization and remodeling of the late ECM, which is subsequently replaced by bone [10]. Notably, terminal hypertrophic differentiation does not occur in healthy articular cartilage [11]. To investigate chondrogenesis, a variety of experimental models have been established. In addition to established cell lines such as ATDC5 [12] or C3H10T1/2 [13], induced pluripotent stem cells (iPSCs) [14] and mesenchymal stem cells (MSCs) [15–17] have been used to study transcriptional dynamics and ECM development during chondrogenesis. Stem-cell based approaches however are limited by donor variability, limited reproducibility, restricted cell availability and replicative senescence [18–20]. Hence, simpler models such as ATDC5 cells offer an advantage of higher robustness, reproducibility and scalability as well as effective gene editing possibilities for comparative studies of genetic variations.

The ATDC5 model system has been widely used to study chondrogenesis since its first description in 1990 by Atsumi et al. [12] who also established the cell line and successfully induced differentiation by supplementing the growth medium with insulin, transferrin and sodium selenite, a protocol which still remains a standard today. Subsequently, additional supplements such as ascorbic acid or β-glycerophosphate have been introduced to accelerate chondrogenic differentiation [21–23] or enhance matrix mineralization [24,25].

Surprisingly, despite more than 1000 PubMed entries related to ATDC5 cells as of January 2026, a comprehensive and unbiased characterization of ATDC5 gene expression patterns during differentiation as well as composition of the secreted ECM are still lacking: most studies using ATDC5 cells have followed the expression of only few chosen chondrocyte marker genes [26] such as *Sox9, Col2a1* and *Col10a1*. Moreover, only a single study [27] has performed an extensive proteomic characterization of ATDC5 ECM.

Here, we combined time-resolved transcriptomic analyses with ECM-focused proteomics to provide an integrated characterization of transcriptional dynamics and matrix composition during ATDC5 chondrogenesis.

## 2. Materials and Methods

### 2.1 Cell culture

ATDC5 cells were obtained from ECACC. Cells were cultured in medium composed of Dulbecco’s modified eagle medium (DMEM, 32430100, Gibco™, Thermo Fisher Scientific, Waltham, MA, USA) and Ham’s F-12 (11765054, Gibco™, Thermo Fisher Scientific, Waltham, MA, USA) in a ratio of 1:1, supplemented with 10 % fetal calf serum (FCS, F7524 Sigma, Taufkirchen, Germany), 1% Sodium Pyruvate (S8636, Sigma, Taufkirchen, Germany), 100 U/ml penicillin and 100 µg/ml streptomycin (P-0781, Sigma, Taufkirchen, Germany) at 37°C and 5% CO_2_. The cells were passed every 3-4 days.

For differentiation experiments, ATDC5 cells were plated at 3×10^5^ or 0.75×10^5^ cells/well into 6 or 12-well plates (83.3920/ 83.392, Sarstedt, Nümbrecht, Germany). Upon confluency, the growth mediums FCS concentration was reduced to 5% and was supplemented with 10µg/ml insulin (I9278, Sigma, Taufkirchen, Germany), 30nM sodium selenite (S5261, Sigma, Taufkirchen, Germany), 50µg/ml ascorbic acid (A4403, Sigma, Taufkirchen, Germany) and 10µg/ml transferrin (90190, Sigma, Taufkirchen, Germany). The medium was changed every other day.

### 2.2 Proteoglycan and collagen staining

Cells were washed twice with PBS (D8537, Sigma, Taufkirchen, Germany) and fixed with ice-cold methanol. At indicated time points, cells were stained for proteoglycans and collagens using 0.1% Alcian Blue (A5268, Sigma, Taufkirchen, Germany) in 0.1M HCl and Sirius Red solution (13422, Morphisto, Offenbach am Main, Germany) for 2 hours at room temperature. Following incubation, excess dye was removed. Alcian Blue-stained cells were rinsed with PBS, while Sirius Red-stained cells were washed with 0.01 M HCl to remove unbound dye. Images of the stained cultures were captured using a Leica microscope or phone camera.

### 2.3 Immunohistochemistry

Cells were seeded on gelatin-coated cover slips and maintained until they reached confluency. Cells were washed three times with PBS, fixed with 4% PFA solution. After blocking with 5% BSA in PBS solution the fixed cells were stained with rabbit anti fibronectin antibody (1:500, ab2413, abcam, Cambridge, United Kingdom) for 2h. Cells were then washed three times with PBS and incubated with Alexa 488 Goat Anti Rabbit (1:500, A11008, Thermo Fisher Scientific, Waltham, MA, USA). Cells were finally washed three times with PBS and mounted in Vectashield with DAPI (VEC-H-1200, Biozol, Hamburg, Germany). Images were taken with a Leica THUNDER Imager (Leica, Wetzlar, Germany) with a 63X objective.

### 2.4 Proliferation assay

Cell proliferation was assessed using the Click-iT™ Plus EdU Alexa Fluor™ 555 Imaging Kit (C10638, Thermo Fisher Scientific, Waltham, MA, USA) following the manufacturer’s instructions. Briefly, cells were seeded on coverslips and incubated with 10 μM EdU (5-ethynyl-2′-deoxyuridine) for 2 hours at the indicated time points. After incubation, cells were fixed with PFA 5% in PBS, permeabilized with Triton 0.5% in PBS, and subjected to the Click-iT™ Plus reaction cocktail for EdU detection. Cover slips were mounted in Vectashield with DAPI. Fluorescent images were acquired at different spots on the cover slip. EdU incorporation was evaluated by analyzing 3 images, but at least 100 nuclei, per time point.

### 2.5 RNA sequencing

RNA was extracted from cells grown in 6-well plates at indicated time points using RNeasy Fibrous Tissue Mini Kit (74704, Qiagen, Hilden, Germany) according to the protocol provided by the manufacturer. RNA concentration was measured with Nano Drop ND-1000 (Thermo Fisher Scientific, Waltham, MA, USA). RNA quality was assured by measurement of RNA integrity value using High Sensitivity RNA Screen Tape Analysis (Tape Station 4150, Part number 5067 5579, Agilent technologies, Santa Clara, CA, USA). RNA sequencing of four replicates per time point was performed at Novogene, Hongkong. For each sample 5 GB raw data were available. All RNA sequencing data generated in this study have been deposited in the Gene Expression Omnibus under accession number GSE324360.

### 2.6 Transcriptomics bioinformatics

RNA-Seq data analysis was performed using the Galaxy web platform (usegalaxy.eu, Galaxy). Raw sequencing reads were first quality-checked using FastQC (version 0.74+galaxy1), and adapter trimming was carried out with Trim Galore! (version 0.6.7+galaxy0). One sample at this time point could not be processed in Galaxy for unknown reasons. As three additional biological replicates were available, this sample was excluded from downstream analysis.

Cleaned reads were then mapped to the mouse reference genome (GRCm39) using RNA STAR (version 2.7.11a+galaxy1) as paired end data and “length of the genomic sequence around annotated junctions” set to 149. Read counts per gene were quantified using featureCounts (version 2.0.6+galaxy0) with GENCODE comprehensive gene annotation for mouse (GRCm39, release M35; Ensembl 112). The resulting count matrix was used as input for differential gene expression analysis in DESeq2 (version 2.11.40.8+galaxy0) within Galaxy. Normalized count files were used for the generation of dot plots encoding gene-wise z-scores by color and scaled values by point size, using the pheatmap package in R. GO term enrichment analysis was conducted using the Functional Annotation tool in DAVID [28] to identify significantly enriched biological processes. GO enrichment analyses were conducted using the complete gene sets, unless when a high proportion of not annotated genes hampered GO analyses in which case not annotated genes were excluded from GO analysis; such cases are indicated in the figure legends. Graphs and Volcano plots were created using Prism10 GraphPad Software (v10.5.0, GraphPad Software, Boston, MA, USA).

For creating the k-means clustered heatmap the results were filtered for adjusted p < 0.05 and log_2_(fold change (FC)) ±2 as indicated in the figure legends. Before fitting and statistical analysis, counts were normalized by library size (DESeq2::estimateSizeFactors) and gene-wise dispersion (DESeq2::estimateDispersions) to normalize the gene counts by gene-wise geometric mean over samples. For plotting the heat maps, gene-wise scaling was performed on the normalized counts to cluster the counts for the expression tendency between the samples (stats::kmeans function). In addition, the counts were centred by setting the gene-wise mean to zero, which explains why some count values are negative (below gene-wise mean). Visualization of the clustered data was performed using the pheatmap::pheatmap function (pheatmap v.1.0.10) with deactivated function-intrinsic clustering.

### 2.7 ECM proteomics

ATDC5 cells were cultured in differentiation medium until harvest. Cells were removed by incubating the sample with decellularization buffer (20 mM NH₄OH, 0.5% Triton X-100), followed by extensive PBS washing and 30 min DNAseI (15U/ml, 79254, Qiagen, Hilden, Germany) treatment. Lysis of remaining ECM and preparation for mass spectrometry analysis was performed as described previously [29]. Peptides were analyzed with the Evosep One system (Evosep Biosystems, Odense, Denmark) coupled to a timsTOF fleX mass spectrometer (Bruker, Billerica, MA, USA). 500 ng of peptides were loaded onto Evotips C18 trap columns (Evosep Biosystems, Odense, Denmark) according to the manufacturer’s protocol. Peptides were separated on an EV1137 performance column (15 cm x 150 µm, 1.5 µm, Evosep, Odense, Denmark) using the standard implemented 30 SPD method with a gradient length of 44 min (buffer A: 0.1% v/v formic acid, dissolved in H_2_O; buffer B: 0.1% v/v formic acid, dissolved in acetonitrile). Over the time of the gradient, the concentration of acetonitrile gradually increased from 0 to 90% at a flow rate of 500 nl/min.

The timsTOF fleX mass spectrometer (Bruker, Billerica, MA, USA) was operated in the DIA-PASEF mode. DIA MS/MS spectra were collected in the range in an m/z range from 100 to 1700. Ion mobility resolution was set to 0.60–1.60 V·s/cm over a ramp time of 100 ms and an accumulation time of 100 ms. The cycle time was set at 1.8s. The collision energy was programmed as a function of ion mobility, following a straight line from 20 eV for 1/K0 of 0.6 to 59 eV for 1/K0 of 1.6. The TIMS elution voltage was linearly calibrated to obtain 1/K0 ratios using three ions from the ESI-L TuningMix (Agilent) (m/z 622, 922, 1222).

Raw data were analyzed with DIA-NN software (v.2.0) [30]. A spectral library was predicted using a FASTA file containing the murine protein sequences as of June 23rd, 2025 (mouse-EBI-reference database, https://www.ebi.ac.uk/). The false discovery rate (FDR) was set to 1%. The search was performed allowing one missed cleavage and cysteine carbamidomethylation enabled as a fixed modification. Match between runs was enabled. Quantification was performed using the label-free quantification algorithm MaxLFQ, which calculates the protein quantities as ratios from all peptide intensities.

The data was further processed and analyzed for the statistical analysis using R (v4.3.0) within R studio. Protein intensities were log2-transformed and median normalized. Differential expression analysis was performed using limma [31]. P-values were corrected by multiple testing using the Benjamini–Hochberg method, as applied by limma. Mass spectrometry raw data have been deposited at the ProteomeXchange Consortium (http://proteomecentral.proteomexchange.org) under the accession number PXD074419. Furthermore, all mass spectrometry proteomics datasets used and/or analysed during this study are available online at the MassIVE repository (http://massive.ucsd.edu/; dataset identifier: MSV000100830; Reviewer account details: Username: “MSV000100830_reviewer”, Password: “ST4796_Schmidts"). This study did not generate new code for analysis.

Dot plots encoding gene-wise z-scores by color and scaled values by point size were generated using the pheatmap package in R. For creation of Venn Diagrams interactivenn web tool was used [32].

### 2.8 Statistical analysis

Data are presented as mean ± SD. Statistical analyses were performed using appropriate tests as indicated in the figure legends. For proliferation analysis two-way ANOVA followed by Sidak’s multiple comparisons test was used to perform comparisons of differentiated to the control cultures. For transcriptomic and proteomic data, p-values were corrected with Benjamini-Hochberg method and adjusted p-values < 0.05 were considered statistically significant.

## 3. Results

### 3.1 Marked reduction of cell proliferation and increased ECM deposition upon ATDC5 cell differentiation

ATDC5 cells have been previously reported to produce a cartilage-like ECM composed of collagens and proteoglycans upon differentiation [22,27,33]. We used a standard protocol to differentiate ATDC5 cells with insulin, transferrin, sodium selenite and ascorbic acid [21] for 4, 7, 14 or 21 days. As expected, visualisation of deposited ECM using Alcian Blue and Sirius Red [22] revealed progressive matrix accumulation over time, more pronounced in cells treated with differentiation medium compared to cells in control medium (Figure 1 A). Immunostaining for fibronectin, a protein binding to collagen fibrils and regulating fibril assembly [34,35] at differentiation day 4 revealed the formation of an unorganized fibrillar ECM network (Figure 1 B). In vivo, cellular differentiation of chondroprogenitor cells into first proliferating, then hypertrophic chondrocytes goes along with an initially increasing proliferation rate, followed by a termination of proliferation once cells reach hypertrophic stages of chondrogenesis [10]. We therefore tested the impact of differentiation on ATDC5 cell proliferation which revealed observed a marked decrease in cell proliferation already by day 4 for both cells treated with control or with differentiation medium. By day 7, cell proliferation was undetectable for cells treated with differentiation medium and only minimal proliferation was observed for control medium cultures (Figure 1 C). Potentially, as ATDC5 cells are derived from teratocarcinoma cells [12] with a fairly high proliferation rate already, increasing cell density over time could impact cell proliferation differently to what is observed upon in vivo chondrocyte differentiation. Other proliferation studies in ATDC5 cells have similarly shown that cell number and proliferation rate decrease over time [23,36,37].

**Figure 1:**
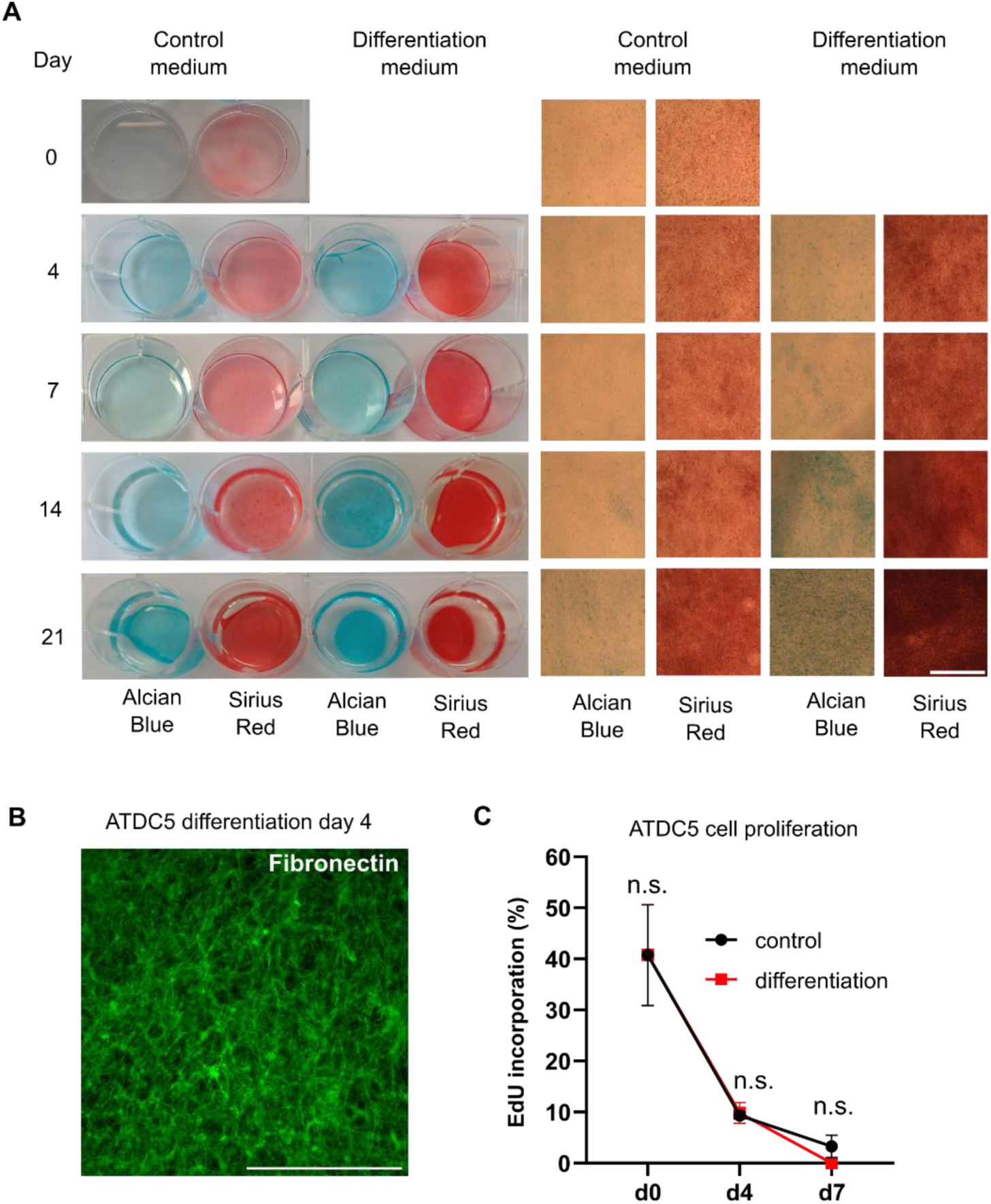
ECM deposition and cell proliferation during ATDC5 differentiation. Cells were maintained in control medium (5% FCS) or differentiation medium (5% FCS, supplemented with 10µg/ml insulin, 30nM sodium selenite, 50µg/ml ascorbic acid and 10µg/ml transferrin). (A) Alcian Blue (marking proteoglycans) and Sirius Red (marking collagens) staining of cells maintained in control or differentiation medium at days 0, 4, 7, 14 and 21 of culture visualising progressive ECM accumulation over time. Scale bar: 1mm. (B) Fibronectin staining of ATDC5 derived ECM after decellularization at day 4 of culture in differentiation medium. Scale bar: 100µm. Images were taken using a Leica Thunder microscope. (C) To assess proliferation rates, the proportion of EdU (5-ethynyl-2′-deoxyuridine) positive nuclei was determined at day 0, 4 and 7 for cells maintained in control or differentiation medium. Data are presented as mean ± SD (n=3). Two-way ANOVA followed by Sidak’s multiple comparisons test was used to perform pairwise comparisons of differentiating cultures compared to control at each time point. P-values < 0.05 were considered statistically significant.

### 3.2 Transcriptional changes reflecting ATDC5 chondrogenic differentiation over time

ATDC5 cells are proposed to undergo a multistep chondrogenic differentiation, encompassing proliferative and hypertrophic stages [24,33,38,39] mimicking the process of endochondral bone formation. However, this conclusion is based mainly on studying expression of few marker genes, while genome-wide studies to investigate effects on the transcriptome are lacking. We therefore performed transcriptome analyses at day 0, 4, 7, 14 and 21 of cells cultured in differentiation medium.

Principal component analysis (PCA) demonstrated distinct clustering of samples at all time points, with samples from day 14 and day 21 showing particularly high similarity (Figure S1). K-means clustering of normalized scaled counts identified five distinct gene clusters (Figure 2 A). We subsequently used GO enrichment analysis to functionally characterize the genes within each cluster and to derive descriptive cluster annotations from most enriched GO terms. Cluster I (Translational initiation and uncharacterized genes), marked by genes with initially high but sharply declining expression after day 0, mainly consisted of predicted or poorly annotated genes. The remaining genes were predominantly associated with the GO term translation initiation (Figure 2 B). Genes in cluster II (Immune response), the smallest cluster, showed a pronounced increase in expression shortly after the onset of differentiation at day 4 and were predominantly associated with the innate immune response (Figure 2 C). Genes in cluster III, (ECM I) showed a modest increase between day 0 and 7 followed by a decrease until day 21 with GO term extracellular matrix organization (GO:0030198) found to be the most significant (Figure 2 D), comprising 13 genes including *Abi3bp* and *Pdgfra*, which have known roles in early chondrogenesis [40,41]. Cluster IV (ECM II), was characterized by peak gene expression at day 14 and showing enrichment for genes related to cartilage development and ossification, encompassing key chondrocyte marker genes [26,42] (*Acan*, *Col2a1*, *Hapln1*) (Figure 2 E). Genes in clusters III and IV, both enriched for ECM and cartilage genes, showed lowest expression before culturing in differentiation medium, indicating that their expression is enhanced by the culture conditions. The largest cluster, cluster V (Angiogenic/metabolic response), showed a continuous increase in gene expression until day 21 (Figure 2 F). GO analysis of all genes in clusters I-V revealed that genes that change most in expression during the differentiation process are associated with ECM and immunity linked GO terms (Figure S2). Gene cluster lists and complete GO enrichment results are provided in Supplementary File S1.

**Figure 2:**
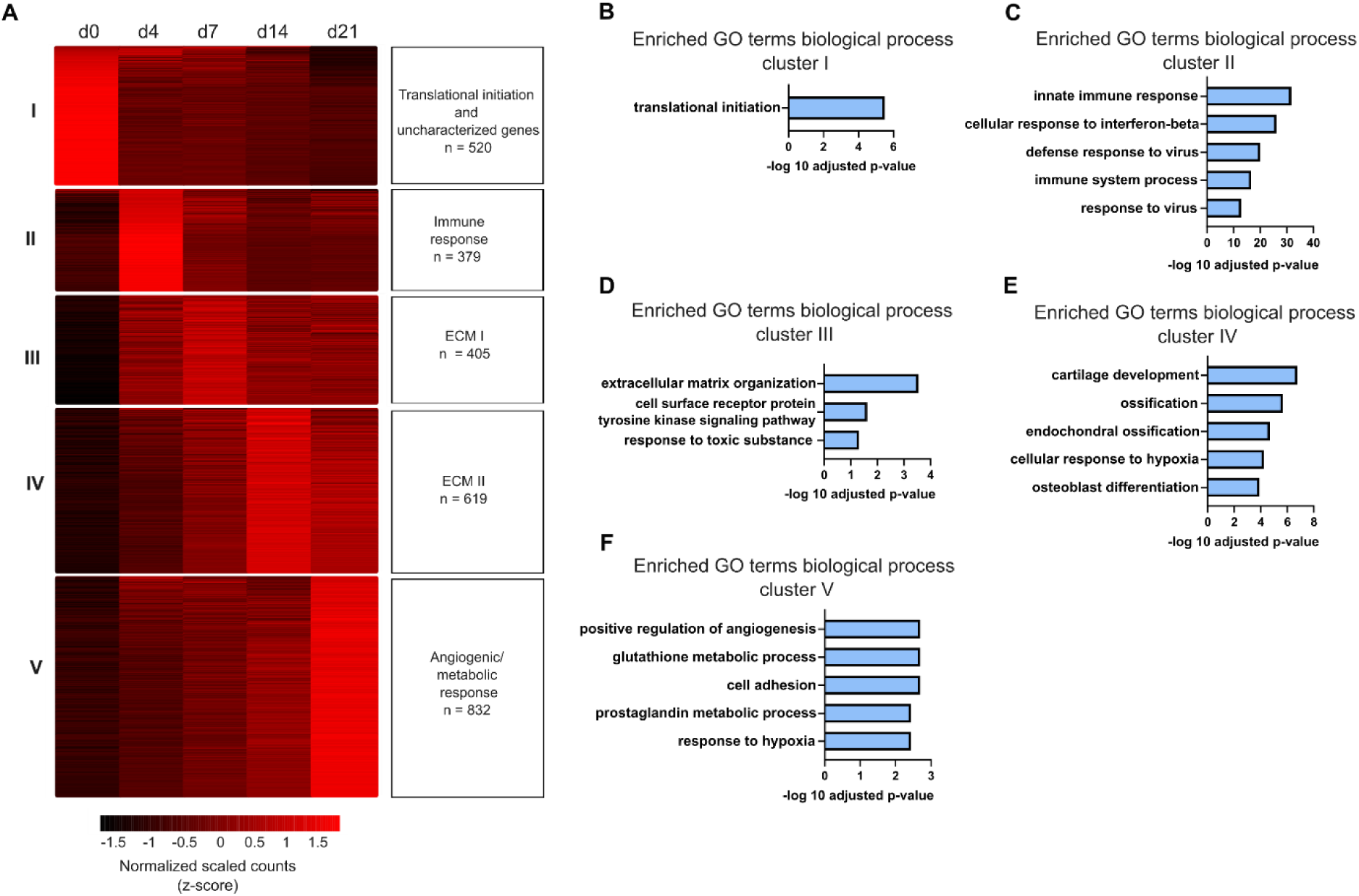
Early immune response followed by ECM organization and lastly and angiogenic/metabolic response expression patterns over the course of ATDC5 differentiation. (A) Differentially expressed genes (adjusted *p* ≤ 0.05, |log2 fold change| ≥ 2) were identified in ATDC5 cells maintained in differentiation medium at days 0, 4, 7, 14, and 21. Differential expression was assessed using DESeq2 based on a two-sided Wald test with Benjamini–Hochberg correction. Genes were grouped into five clusters using k-means clustering. The heat map displays z-score scaled normalized expression values. (B-F) Gene ontology (GO) analysis for “biological process” (BP) of all genes in clusters I-V using DAVID Functional Annotation Tool. Top 5 or all identified GO terms are displayed. For analysis of cluster I (B), genes that were no annotated (predicted genes) were excluded. Full list of genes and GO analysis per cluster are listed in Supplementary File S1.

To complement the clustering of normalized expression profiles, we further analyzed gene expression changes between consecutive sampling time points (Figure 3 A-D, Supplementary File S2). When comparing the next later sampling timepoint to the previous sampling timepoint (day 4 to day 0, day 7 to day 4, day 14 to day 7 and day 21 to day 14), the highest number of differentially expressed genes was found between day 0 and day 4 (Figure 3 A, E). As differentiation progressed, the number of differentially expressed genes between measurements decreased (Figure 3 E). Comparison of day 4, day 7, day 14 and day 21 to day 0 directly revealed significantly increased expression of key chondrocyte genes such as *Col2a1*, *Acan*, and *Fmod* and the GO term “Extracellular Matrix organization” among the three most significant at all time points later than day 4 (Figure 3 A, Figure S3, Figure S4, Supplementary File S3). While expression of these and other chondrocyte markers already increased between day 0 and day 4 (Figure 3 A), the majority of genes showing increased expression at day 4 were not related to chondrocyte differentiation but rather associated with immune response as revealed by GO analysis (Figure S5 B, Supplementary File S3). Concurrently, genes related to cell division decreased in expression (Figure S5 A). The expression of chondrocyte differentiation, ECM- and ossification-related genes increased over time, most prominently between day 7 and day 14, as exemplified by increased expression of *Col10a1*, *Hapln1*, and *Comp* (Figure 3 C), together with strong enrichment of GO terms related to cartilage development and ossification (Figure S5).

**Figure 3:**
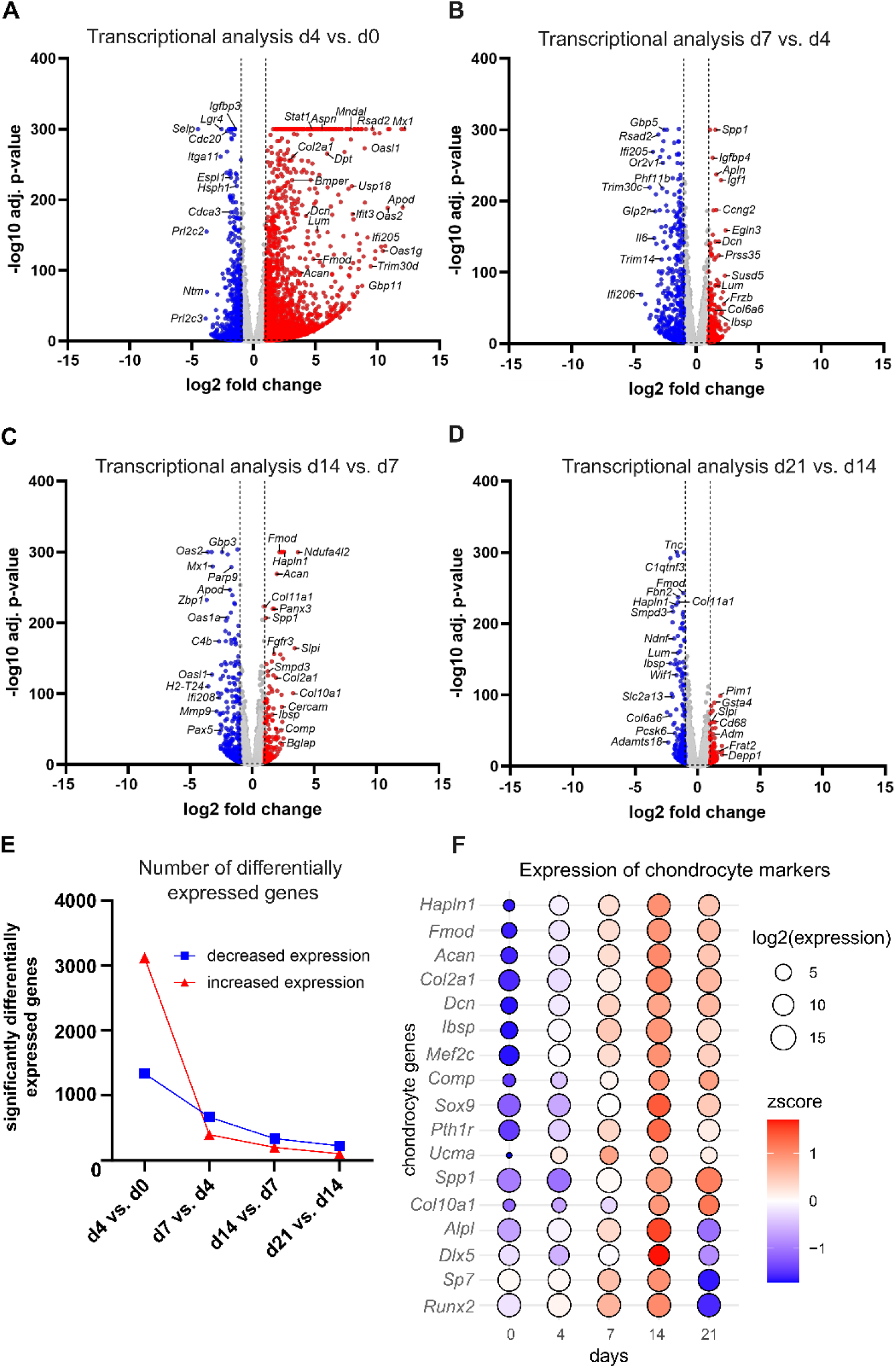
Increasing expression of key chondrocyte marker genes upon differentiation peaking at day 14. (A-D) Volcano plots display differential gene expression between consecutive time points based on RNA-seq data, revealing significant increase in expression of chondrocyte differentiation genes such as *Col2a1*, *Acan* and *Fmod*. The x-axis indicates log2 fold change, the y-axis shows the −log10 adjusted p-value. For visualization purposes, −log10 adjusted p-values were capped at 300. Genes were color-coded according to log2fold change thresholds of ±1. (E) Number of differentially expressed genes with increasing or decreasing expression during sampling intervals, showing overall decreasing numbers of differentially expressed genes as differentiation progressed. (F) Dot plot displaying the expression of selected chondrocyte marker genes across the indicated time points with peak expression at day 14. Dot size represents log2-transformed expression values, while color intensity indicates z-score-scaled expression levels for each gene. Expression values are derived from DESeq2-normalized RNA-seq counts.

Representative chondrocyte differentiation and ECM genes [43,44] including collagens, proteoglycans, and transcription factors, showed a gradual increase in expression from day 0, peaking at day 14 (Figure 3 F) with only few exceptions. *Col2a1*, a marker of the proliferative chondrocyte phase [45], was representative of this general expression pattern (Figure S1 B). In contrast, expression of *Col10a1*, a marker of hypertrophic differentiation [46], increased only modestly during early differentiation but rose sharply from day 7 onwards and continued to increase beyond day 14 (Figure S1 C). Similarly, *Spp1*, encoding the osteogenic glycoprotein osteopontin involved in bone mineralization [47], reached its highest expression level at day 21. *Ucma*, encoding a negative regulator of osteogenesis usually expressed in early chondrogenesis [48], showed highest gene expression at day 7. Interestingly, *Sox9* and *Runx2*, master transcription factors of chondrogenic [49,50] and osteogenic differentiation [51], reached their highest expression levels at day 14, although their overall expression increased only modestly over the course of differentiation (Figure 3 F, Figure S1 D). The overall stable expression of Sox9 is in accordance with previous ATDC5 studies [21,23,37,52,53], while for Runx2, studies have reported either a progressive increase in expression over time [21,23] or a transient increase followed by a decline [37,53]. Published in vivo data suggests that Sox9 expression is typically declining during hypertrophic differentiation, however Runx2 expression has been reported to increase upon chondrocyte prehypertrophy [10,54,55].

When comparing the intervals between day 7 vs. day 4, as well as between day 14 vs. day 7, immunity-related genes showed a decrease in expression, as indicated by GO analysis (Figure S5 C, E), consistent with the expression pattern observed for cluster II, which could indicate a stress effect resulting from addition of differentiation medium and cells reaching full confluency.

### 3.3 Dissection of the ATDC5 derived cartilage-like proteome

To investigate the composition of the cartilage-like matrix generated by ATDC5 cells over time upon differentiation, we performed quantitative proteomics analyses. Samples collected after 4, 7, 14 or 21 days of differentiation were decellularized while on day 0, cells were retained.

Principal component analysis revealed clear clustering of samples by time points (Figure 4 B), with day 0 samples clearly separated from all other timepoints. In addition, while day 4, 7 and 14 samples clustered close together, day 21 samples appeared distinct although comparatively few ECM proteins showed differential expression between day 14 and day 21 (Figure 4 F).

**Figure 4:**
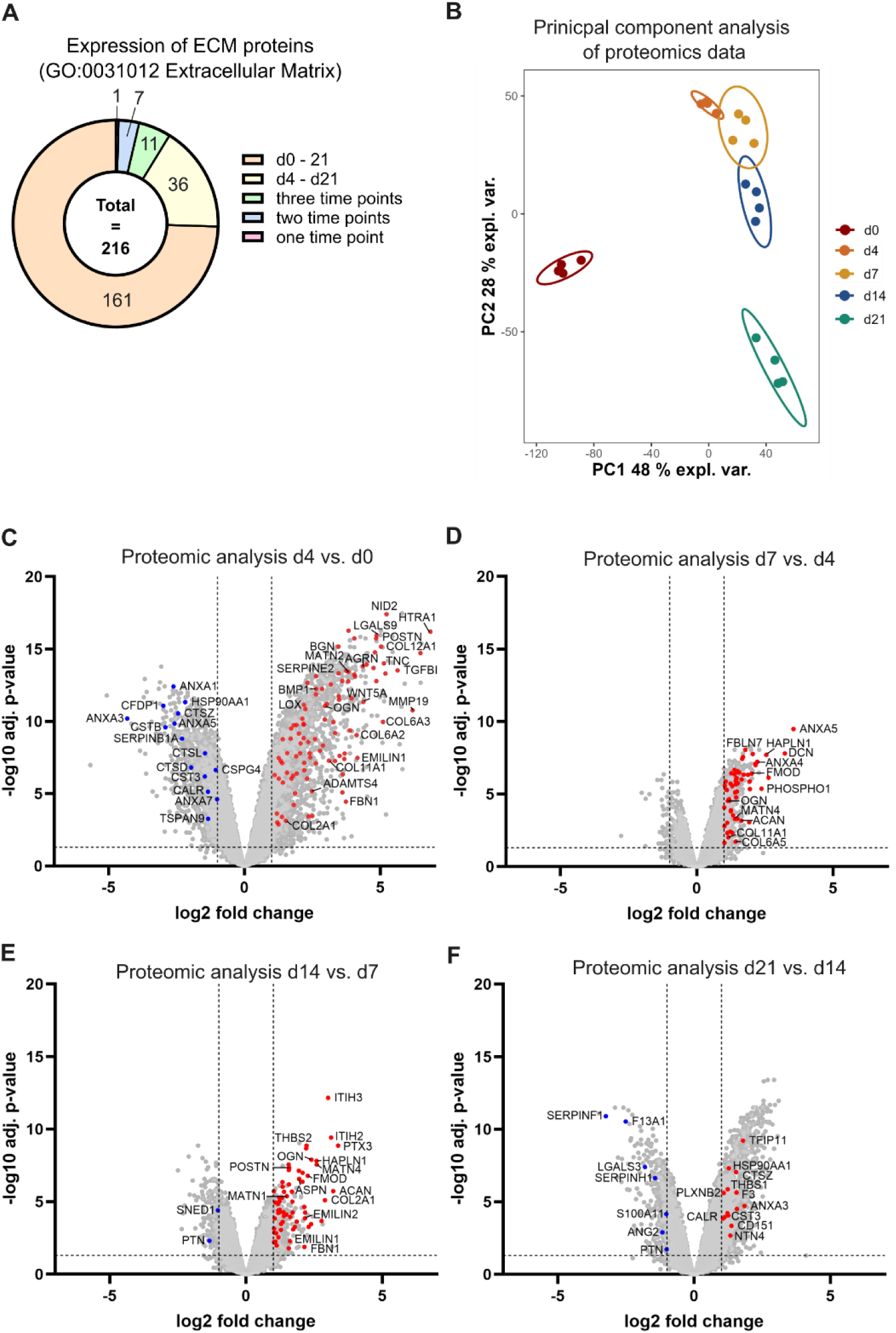
Proteomics analyses reveal increasing expression of ECM proteins ATDC5 derived cartilage-like organoids over time. (A) Donut plot illustrating the number of ECM proteins, annotated in GO:0031012 ECM, that were detected throughout the experiment, at day 4, 7, 14 and 21, or at three or fewer time points. A total of 161 proteins were consistently detected at all time points, whereas 36 proteins were detected from day 4 onwards (including day 4) but not at day 0. Only few proteins were detected at three or fewer time points. For a complete list of all identified proteins and the time points at which they were detected, refer to Table 1. (B) Principal component analysis (PCA) of proteomic samples based on log₂-transformed and median-normalized protein intensities, showing clustering of samples by time point. PC1 and PC2 explain 48% and 28% of total variance. Samples are color-coded according to differentiation time (day 0, 4, 7,14 or 21). (C-F) Volcano plots display differential protein expression between consecutive time points, illustrating a progressive increase in ECM protein expression at each successive time interval. ECM proteins were color-coded according to adjusted p≤0.05 and log2fold change thresholds of ±1. The x-axis indicates log2 fold change, the y-axis shows the −log10 adjusted p-value. Proteins not annotated in GO:0031012 ECM are displayed in grey.

**Table 1:**
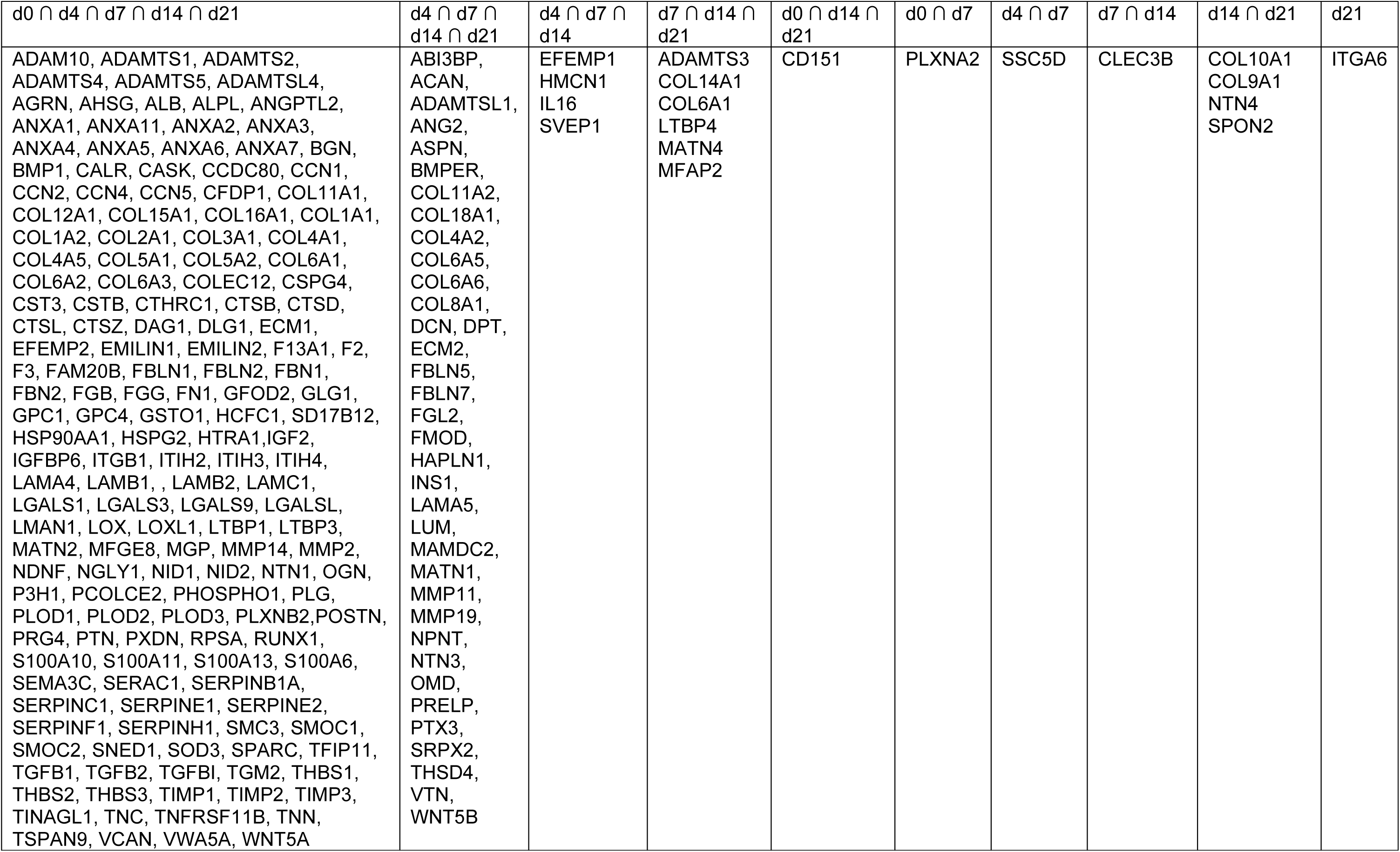
ECM proteins detected in ATDC5 derived cartilage-like organoids. Proteins annotated to GO:0031012 (Extracellular matrix) identified in this study and ordered by detection time points

A protein was considered identified if it was detected in at least two out of four biological replicates at any time point. This led to identification of 8316 proteins across all samples, further referred to as the ATDC5 proteome (Supplementary file S4). Of those, 216 proteins were part of the Gene Ontology term “extracellular matrix” (GO:0031012) (Table 1) of which 161 proteins (74.5%) were consistently identified at all time points (Figure 4 A), including collagens (COL2A1, COL6A1, COL11A1), proteoglycans (CSPG4, BGN, HSPG2), glycoproteins (EMILIN1, MATN2, SMOC1) and ECM-modifying enzymes (ADAMTS5, HTRA1, LOX). 36 proteins were expressed at all time points between day 4 and day 21 after differentiation onset but not day 0, indicating that their expression is induced by the culturing conditions (Figure 4 A). These included aggrecan core protein (ACAN), forming the core of the most abundant proteoglycan in cartilage [56,57], and hyaluronan and proteoglycan link protein 1 (HAPLN1), a key structural component of the cartilage matrix [58]. Several other collagens and proteoglycans were also newly detected during this period including ASPN, DCN, FMOD and LUM, in accordance with their increased expression on gene level comparing day 4 vs. day 0 (Figure 3 A). 19 of 216 ECM proteins were detected only at three or fewer time points (Figure 4 A, Table 1). Beyond day 4, expression of proteins annotated in GO:0031012 “extracellular matrix”, increased almost exclusively over time, with very few exceptions (Figure 4 D-F). Comparison of ECM protein expression at days 4, 7, 14 and 21 relative to day 0 revealed a consistent expression pattern, with a high number of proteins increasing in expression and a small subset of proteins showing lower abundance over time, including cathepsin proteases (CTSZ, CTSB, CTSL, CTSD), which are found intra- and extracellularly [59]. Across all time points, HTRA1, POSTN, BGN, NID2, COL12A1 and MMP19 were among the most significant increasingly expressed ECM proteins (Figure 4 C, Figure S6). Notably, COL10A1, was only identified at day 14 and day 21, consistent with its pronounced increase in gene expression observed at day 14 in transcriptomic analysis (Figure S1 C).

To assess whether ECM protein expression followed specific temporal patterns, representative proteins were visualized in heatmaps (Figure 5). The majority of proteins reached peak expression at either day 14 or day 21, consistent of a progressive accumulation of secreted proteins over time. Proteins were grouped into functional categories: collagens, proteoglycans, glycoproteins, ECM receptors, and ECM-modifying enzymes, thereby encompassing the most relevant ECM components. Collectively, the heatmaps show that the majority of ECM proteins increase steadily over time, independent of functional category. For a comprehensive heatmap including all 216 ECM proteins identified in our dataset see Figure S7.

**Figure 5:**
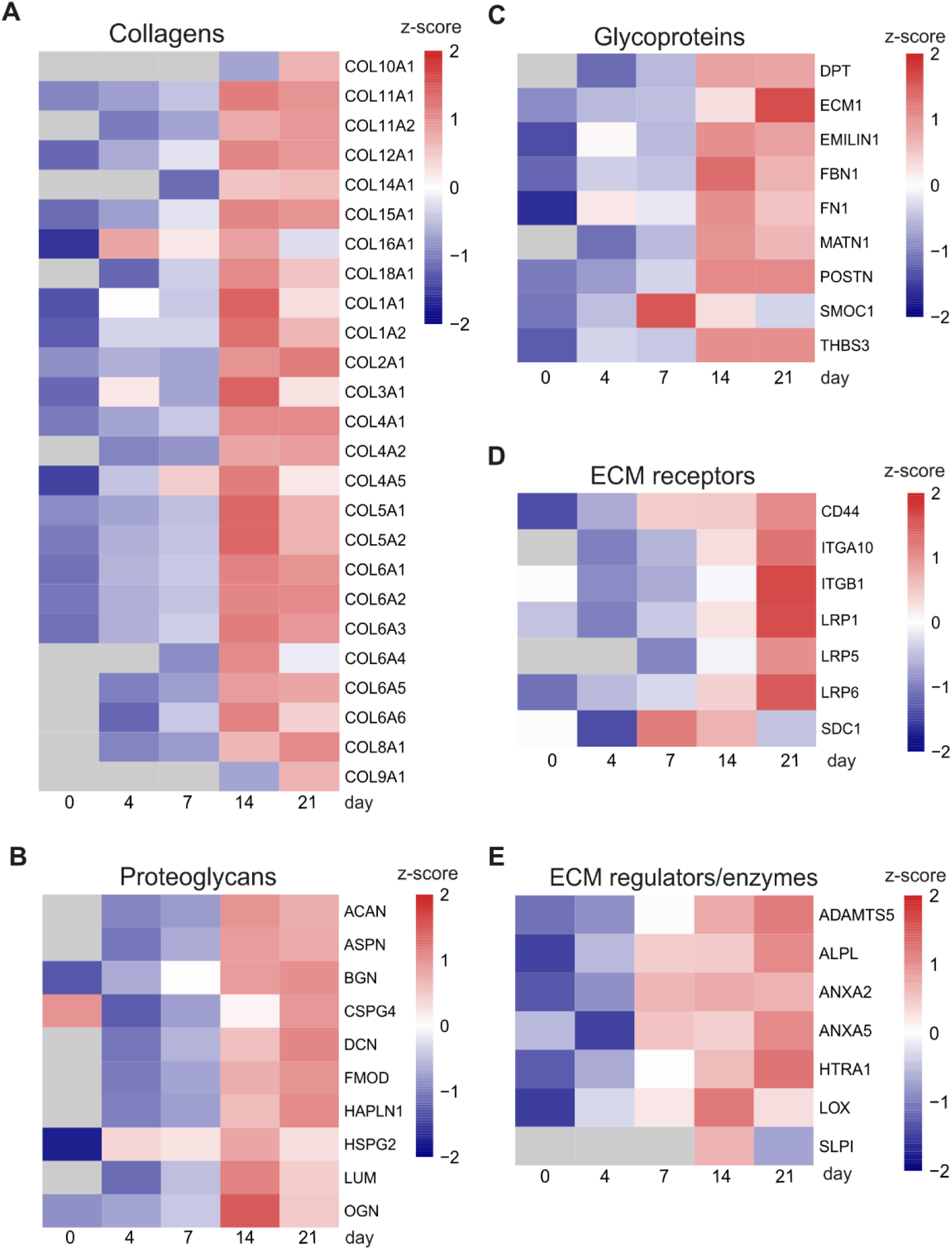
Consistent increase in ECM protein abundance across functional categories during ATDC5 cell differentiation. (A-E) Heatmaps of representative ECM proteins across functional categories (collagens, proteoglycans, Glycoproteins, ECM receptors, ECM regulators/enzymes). Most proteins show low expression at the beginning of differentiation, with peak expression observed at day 14 or 21, with only a few exceptions. Selected proteins from different ECM categories are shown, with normalized values derived from the log2 intensities (expression matrix). Expression values are scaled per protein (z-score) to illustrate relative abundance patterns across timepoints.

### 3.4 Concordant temporal patterns in transcriptomic and proteomic analyses

In general, transcriptomic and proteomic analyses revealed similar temporal trends. In both datasets, ECM-associated genes and ECM proteins increased in expression following the onset of differentiation, with strong induction already observed at day 4 compared to day 0 (Figure 3 A, Figure 4 C). In both analyses, the main increase in ECM genes or proteins occurred between day 7 and 14. In the transcriptomic analysis, expression levels of ECM related genes declined again after day 14, while in the proteomic dataset only few ECM proteins showed significant decreases or increases in abundance between day 14 and 21, indicating limited changes in ECM protein abundances despite changes in gene expression.

### 3.5 ATDC5-derived ECM contains a broader spectrum of ECM proteins than previously reported

ECM proteome composition over time during ATDC5 differentiation has been unclear prior to this study. Therefore, we aimed to systematically investigate the ATDC5-derived ECM proteome and place it in the context of the only other existing dataset previously published by Wilhelm et al. [27]. To do this, we used the Gene Ontology terms “extracellular matrix” (GO:0031012) as well as the Matrisome database [60] as references, as those provides a well-established and widely accepted framework for the functional classification of genes and proteins and compared our proteomic dataset to Wilhelm et al [27].

This revealed 62 ECM proteins commonly detected in our study as well as by Wilhelm et al [27] (Figure 6 A, Supplementary File S5), including the highly abundant fibrillar collagen COL2A1, as well as less abundant collagens including COL6A1, COL6A2, COL11A1, COL11A2 and COL10A1 [61], the main cartilage proteoglycan aggrecan as well as several small leucine-rich proteoglycans (SLRPs), glycoproteins and ECM-modifying enzymes, reflecting the structural diversity of the cartilage ECM captured in both datasets. Conversely, we could not detect six ECM proteins reported by Wilhelm et al. (COL20A1, COL23A1, COL27A1, COL4A3 and COL4A4). Notably, we detected 154 ECM proteins not previously reported by Wilhelm et al. (Figure 6 A, Supplementary File S5). These proteins spanned a wide range of ECM categories, including minor collagens, proteoglycans, annexins, matrilins, laminin chains, microfibril-associated components, secreted signalling proteins and ECM-remodeling enzymes such as ADAMTS family members and matrix-metalloproteinases (MMPs).

**Figure 6:**
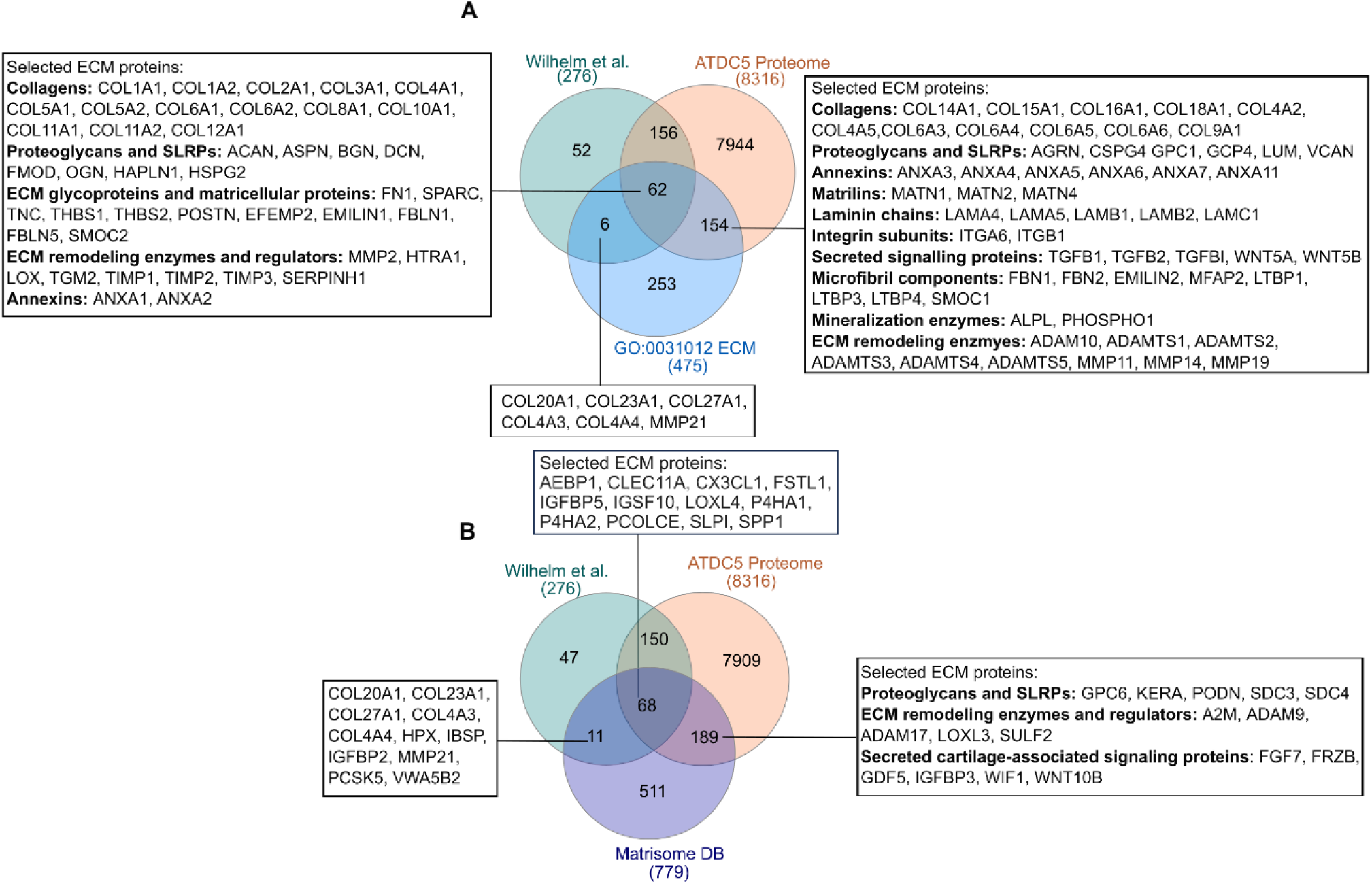
Comparison of detected proteins reveals previously not reported ECM proteins in ATDC5 derived ECM. (A) Venn diagram illustrating overlap of proteins identified in our study (ATDC5 proteome), Wilhelm et al’s study and the GO term “extracellular matrix” (GO:0031012), revealing 154 ECM proteins from numerous functional groups, that were not previously detected in ATDC5 cells (B) Venn diagram illustrating overlap of proteins identified in our study (ATDC5 proteome), Wilhelm et al’s study and the Matrisome database, uncovering 189 ECM proteins not previously reported in ATDC5 cells

In addition, we compared our and Wilhelm et al’s dataset to a list of murine cartilage proteins annotated in the Matrisome Database [60], which is derived from two independent proteomic studies of murine cartilage [62,63] (Figure 6 B). Of 68 proteins that were detected in both our and Wilhelm et al’s study and are listed in the Matrisome Database, 12 proteins were not annotated in GO:0031012 “ECM” (AEBP1, CLEC11A, CX3CL1, FSTL1, IGFBP5, IGSF10, LOXL4, P4HA1, P4HA2, PCOLCE, SLPI, SPP1) (Figure 6 B, Figure S8 A, Supplementary File S5). Further, this comparison revealed 189 proteins only identified in our dataset but not Wilhelm et al.’s study that were also detected in native cartilage, and further 68 proteins that were detected in both our study and by Wilhelm et al (Figure 6 B, Supplementary File S5). Among these 189 uniquely identified proteins, those that were not annotated in GO term “extracellular matrix”, encompassed multiple ECM classes including proteoglycans, ECM remodeling enzymes and regulators, as well as secreted signalling factors (Figure 6 B, Figure S8 A). Comparison of Wilhelm et al.’s dataset with the Matrisome database identified five further ECM proteins (HPX, IBSP, IGFBP2, PCSK5, VWA5B2) that were not detected in our dataset (Figure S8 A). GO analysis of proteins that were neither identified in our nor Wilhelm et al.’s dataset but listed in GO:0031012 or Matrisome database (593 proteins, Figure S8 A) revealed predominantly proteins associated with GO terms “ECM organization”, “proteolysis” and “collagen catabolic process” (Figure S8 B, Supplementary File S6). Overall, our analysis has revealed an extended set of ECM proteins in ATDC5 produced matrix, capturing components that were not previously reported and highlighting the ECM complexity ATDC5-derived cartilage like organoids.

## 4. Discussion

The ATDC5 model system is widely used to study chondrogenic differentiation. Most previous studies relied on targeted approaches to monitor chondrocyte differentiation [22,24,38,64,65], such as qPCR of a low number of selected chondrocyte marker genes, while only very few studies used comprehensive RNAseq [52,66] or ECM proteomic [27] approaches. Thus, there was need for an unbiased genome-wide characterization of gene expression amongst different timepoints ATDC5 in vitro differentiation as well as characterization of the produced ECM. In this study, we combined time-resolved RNA sequencing with ECM proteomics to provide an integrated view of transcriptional dynamics and matrix deposition during ATDC5 chondrogenesis.

Consistent with previous studies [22,24,33,67], staining with Alcian Blue and Sirius Red revealed progressive extracellular matrix deposition over time. Interestingly, proliferative activity decreased rapidly after day 0, when cells were confluent, similarly in both control and differentiating ATDC5 cultures. This shows a limitation of the model as in vivo differentiating chondrocytes are generally assumed to pass through a proliferative phase during early chondrogenesis, characterized by increased proliferative activity under physiological conditions [10]. Our observation could reflect the teratocarcinoma origin of ATDC5 cells resulting in a very high cell proliferation rate at the undifferentiated stage that does not further increase upon differentiation towards chondrogenic cells. Decrease of proliferation of both cells in control and differentiation medium points against any effect linked to differentiation, but could rather reflect effects of an overly dense cell culture after day 0, when full confluency is already reached [68].

At the structural level, in agreement with previously published TEM images of ATDC5 derived ECM [24], fibronectin staining indicated a largely unorganized fibrillar ECM in ATDC5 cultures, which may reflect the constraints of a monolayer culture system and contrasts with the mostly ordered collagen architecture found in native cartilage [69,70]. This highlights an important difference between ATDC5 derived cartilage-like organoids and in vivo cartilage.

Apart from the study by Chen et al. [37], which analyzed the expression of 104 selected chondrocyte-associated genes using real-time RT-PCR, most transcriptional investigations of ATDC5 cells have most commonly focussed on *Col2a1*, *Col10a1* and *Sox9*. Consistent with these reports [21,23,36,37,52,53], our data confirm a strong and early induction of *Col2a1* expression, delayed expression of *Col10a1* and comparatively stable expression levels of *Sox9* and *Runx2* over the course of differentiation. This is of note, as SOX9 is highly, but not exclusively [43,71], expressed in proliferative chondrocytes [45,54], whereas RUNX2 expression is linked to hypertrophic chondrocyte differentiation [55]. This indicates a cell identity change towards a chondrogenic pattern over time, however ATDC5 in vitro differentiation stages do not exactly match known chondrocyte differentiation stages in vivo.

Beyond selected differentiation markers, unbiased transcriptomic analysis revealed that a broad set of chondrocyte- and ECM-related genes increased continuously over time until day 14. Additionally, few genes such as *Col10a1* and *Spp1*, showed peak expression at day 21, demonstrating that ATDC5 cells may have reached a hypertrophic-like stage at this later time points.

While overall, the model seems to recapitulate a key temporal aspect of chondrocyte differentiation, one might have expected clearer separation of ECM gene clusters corresponding to distinct phases of chondrogenesis [72,73]. However, a detailed division between proliferative and hypertrophic stages is not apparent in our data. Potentially, this could be better resolved by investigating more time points. Heterogeneity in the differentiation stage of individual cells, as reflected by nodule formation [38] could also play a role.

Moreover, we identified an immunity cluster with peak gene expression at day 4 and likewise, Farcasanu et al. [66] identified immune-associated Gene Ontology terms in their ATDC5 transcriptomic analysis, using the same differentiation protocol as in our study. This could represent a transient stress response to the changed culture conditions, for example in response to insulin [74] or cell-intrinsic stress responses induced by metabolic constraints and hypoxia in overconfluent cultures [75,76]. However, although not classically associated with chondrogenesis, the activation of innate immune–related gene programs has also rarely been reported during human chondrogenesis [77,78] so that the biological significance of the activation of immune-related genes during chondrogenesis remains to be determined.

Further, the subsequent decline in ECM-related gene expression after day 14, coinciding with increased expression of genes associated with angiogenesis, hypoxia and glutathione metabolism, suggests the activation of transcriptional programs associated with increasingly hypoxic and metabolically stressed conditions secondary to overly dense cell cultures and lack of blood vessels to supply the increasingly thick tissue present in in vivo [79]. This could reflect a major limitation of this in vitro system when investigating late-stage differentiation.

Extracellular matrix makes up the major part of cartilage, providing the environment for differentiating chondrocytes. ECM composition is influenced by chondrocyte differentiation and health but vice versa, ECM may also influence chondrocyte differentiation and homeostasis via matrix-chondrocyte interactions [6]. Hence, understanding the to date largely unresolved composition of ECM secreted by ATDC5 cells is of great interest. Here, we demonstrate that ATDC5 cells produce a more complex ECM than previously appreciated. Of note, Wilhelm et al. applied a detailed fractionation protocol removing the majority of intracellular proteins but also risking losing some ECM components while we only performed decellularization using a buffer composed of 20 mM NH₄OH and 0.5% Triton X-100. To exclude possibly retained intracellular proteins form our conclusions, we chose known ECM and matrisome proteins only to define ATDC5 components. Notably, the identified ECM proteins span all major functional categories, including structural components (collagens, proteoglycans), adaptor proteins (link proteins, matrilins, thrombospondins), surface receptors (integrins, CD44) and various ECM-modifying enzymes, indicating that ATDC5 cells recapitulate multiple functional aspects of cartilage matrix biology. In our study, we additionally identified several collagens that were not previously reported, including minor collagens and fibril-associated Collagen IX, which co-localizes with Collagen II and X to stabilize the fibrillar network [61]. Matrilins (MATN1, MATN2, MATN4) were also detected, acting as adaptors between collagen fibrils and non-collagenous proteins [80]. Our protocol further allowed the identification of numerous secreted components, especially ECM remodeling enzymes, mineralization enzymes and secreted signalling proteins, suggesting ongoing matrix remodelling, calcification and extracellular signalling activity.

Interestingly, neither we nor Willhelm et al. detected COMP, a key ECM protein with roles in promoting chondrogenesis and interacting with several ECM components [81], despite observations of increasing gene expression during differentiation in our study (Figure 3 F). Other protein classes that were not well covered by our and Wilhelm et al.’s dataset comprised predominantly ECM enzymes involved in collagen catabolism and proteolysis. Several of them belong to the family of MMPs, ADAM (A Disintegrin and Metalloproteinase) or ADAMTS (A Disintegrin and Metalloproteinase with Thrombospondin motifs) proteins, which also have implications in osteoarthritis [82]. While we detected some MMPs in our study (MMP11, MMP14, MMP19), we also missed a number, possibly due to low abundance.

Overall, consistent with the transcriptional increase of ECM and cartilage genes over time, ECM protein abundance also accumulated with ongoing differentiation, supporting the biological consistency of the model. In transcriptomic PCA, day 14 and day 21 samples clustered closely, indicating that the overall transcriptional state had not changed dramatically between these time points. In proteomics, however, day 21 samples appeared distinct from day 14, even though relatively few ECM proteins changed in abundance (Figure 4 F), suggesting that intracellular proteins contribute to this separation. Consistent with transcriptomic analyses, the increased gene expression of *Aspn*, *Acan*, *Dcn*, *Fmod* and *Lum* at day 4 compared to day 0 corresponded with the detection of their proteins at day 4, which were not observed at earlier time points. Also, *Col10a1* gene expression increased markedly between day 7 and day 14 and was first detected at the protein level from day 14 onwards. These concordances underscore the consistency of our multi-omics approach. Taken together, these data indicate that under our experimental conditions, chondrogenic differentiation in ATDC5 cells is largely complete between day 14 and 21, consistent with the observed expression of Collagen X and decreased expression of major cartilage genes on day 21.

In summary, our time-resolved transcriptomic and ECM proteomic analysis reveals that ATDC5 cells recapitulate key molecular and matrix-associated features of chondrogenic differentiation. While the system does not fully resolve discrete differentiation stages or reproduce the ordered architecture of native cartilage, it robustly captures major transcriptional programs and produces a complex cartilage-like extracellular matrix. Collectively, these data further establish ATDC5 cells as a robust, accessible and well-characterized platform for mechanistic studies of chondrogenesis and cartilage ECM biology.

## CRediT authorship contribution statement

**Anna Klawonn:** Writing – review & editing, Writing – original draft, Investigation, Formal analysis, Visualization. **Stefan Tholen:** Formal analysis, Investigation. **Ilona Skatulla:** Investigation. **Chiara M. Schröder:** Formal analysis, Visualization. **Sebastian J. Arnold:** Resources, Funding acquisition. **Oliver Schilling:** Resources, Funding acquisition. **Miriam Schmidts:** Writing – review & editing, Writing – original draft, Resources, Project administration, Funding acquisition, Conceptualization. All authors read and approved of the current manuscript.

## Data availability statement

RNA-seq data generated in this study have been deposited in the Gene Expression Omnibus (https://www.ncbi.nlm.nih.gov/geo/) under accession number GSE324360. Mass spectrometry raw data have been deposited at the ProteomeXchange Consortium (https://www.proteomexchange.org) under the accession number MSV000100830).

## Acknowledgement

We thank Christoph Schell and Maximilian Wess for helpful discussion of fibronectin immunofluorescence. The Proteomic Platform – Core Facility (ProtCF) was supported by the Medical Faculty of the University of Freiburg to Prof. Dr. Oliver Schilling (2021/A3-Sch and 2023/A3-Sch). We acknowledge the support of the Freiburg Galaxy Team, funded by the German Federal Ministry of Education and Research [BMFTR](https://www.bmftr.bund.de/DE/Home/home_node.html) grant 031 A538A [de.NBI](https://www.denbi.de/)-RBC and the Ministry of Science, Research and the Arts Baden-Württemberg (MWK) within the framework of LIBIS/de.NBI Freiburg.

## References

1. Bačenková D, Trebuňová M, Demeterová J, Živčák J. Human Chondrocytes, Metabolism of Articular Cartilage, and Strategies for Application to Tissue Engineering. Int J Mol Sci. 2023 Jan;24(23):17096. doi:10.3390/ijms242317096

2. Han L, Grodzinsky AJ, Ortiz C. Nanomechanics of the Cartilage Extracellular Matrix. Annu Rev Mater Res. 2011 Jul 1;41:133–68. doi:10.1146/annurev-matsci-062910-100431 PubMed PMID: 22792042; PubMed Central PMCID: PMC3392687.

3. Hsueh MF, Khabut A, Kjellström S, Önnerfjord P, Kraus VB. Elucidating the Molecular Composition of Cartilage by Proteomics. J Proteome Res. 2016 Feb 5;15(2):374–88. doi:10.1021/acs.jproteome.5b00946 PubMed PMID: 26632656; PubMed Central PMCID: PMC4917199.

4. Svensson RB, Hassenkam T, Grant CA, Magnusson SP. Tensile Properties of Human Collagen Fibrils and Fascicles Are Insensitive to Environmental Salts. Biophys J. 2010 Dec 15;99(12):4020–7. doi:10.1016/j.bpj.2010.11.018 PubMed PMID: 21156145; PubMed Central PMCID: PMC3000511.

5. Farach-Carson MC, Wu D, França CM. Proteoglycans in Mechanobiology of Tissues and Organs: Normal Functions and Mechanopathology. Proteoglycan Res. 2024;2(2):e21. doi:10.1002/pgr2.21 PubMed PMID: 39584146; PubMed Central PMCID: PMC11584024.

6. Gao Y, Liu S, Huang J, Guo W, Chen J, Zhang L, et al. The ECM-Cell Interaction of Cartilage Extracellular Matrix on Chondrocytes. BioMed Res Int. 2014;2014:648459. doi:10.1155/2014/648459 PubMed PMID: 24959581; PubMed Central PMCID: PMC4052144.

7. Grässel S, Aszódi A, editors. Cartilage [Internet]. Cham: Springer International Publishing; 2016 [cited 2026 Jan 24]. Available from: http://link.springer.com/10.1007/978-3-319-29568-8 doi:10.1007/978-3-319-29568-8

8. Unger S, Ferreira CR, Mortier GR, Ali H, Bertola DR, Calder A, et al. Nosology of genetic skeletal disorders: 2023 revision. Am J Med Genet A. 2023 May;191(5):1164–209. doi:10.1002/ajmg.a.63132 PubMed PMID: 36779427; PubMed Central PMCID: PMC10081954.

9. Welsh BL, Sikder P. Advancements in Cartilage Tissue Engineering: A Focused Review. J Biomed Mater Res B Appl Biomater. 2025;113(1):e35520. doi:10.1002/jbm.b.35520

10. Goldring MB, Tsuchimochi K, Ijiri K. The control of chondrogenesis. J Cell Biochem. 2006 Jan 1;97(1):33–44. doi:10.1002/jcb.20652 PubMed PMID: 16215986.

11. Chawla S, Mainardi A, Majumder N, Dönges L, Kumar B, Occhetta P, et al. Chondrocyte Hypertrophy in Osteoarthritis: Mechanistic Studies and Models for the Identification of New Therapeutic Strategies. Cells. 2022 Dec 13;11(24):4034. doi:10.3390/cells11244034 PubMed PMID: 36552796; PubMed Central PMCID: PMC9777397.

12. Atsumi T, Miwa Y, Kimata K, Ikawa Y. A chondrogenic cell line derived from a differentiating culture of AT805 teratocarcinoma cells. Cell Differ Dev Off J Int Soc Dev Biol. 1990 May;30(2):109–16. doi:10.1016/0922-3371(90)90079-c PubMed PMID: 2201423.

13. Shea CM, Edgar CM, Einhorn TA, Gerstenfeld LC. BMP treatment of C3H10T1/2 mesenchymal stem cells induces both chondrogenesis and osteogenesis. J Cell Biochem. 2003;90(6):1112–27. doi:10.1002/jcb.10734

14. Wu CL, Dicks A, Steward N, Tang R, Katz DB, Choi YR, et al. Single cell transcriptomic analysis of human pluripotent stem cell chondrogenesis. Nat Commun. 2021 Jan 13;12:362. doi:10.1038/s41467-020-20598-y PubMed PMID: 33441552; PubMed Central PMCID: PMC7806634.

15. Huynh NPT, Zhang B, Guilak F. High-depth transcriptomic profiling reveals the temporal gene signature of human mesenchymal stem cells during chondrogenesis. FASEB J. 2019 Jan;33(1):358–72. doi:10.1096/fj.201800534R PubMed PMID: 29985644; PubMed Central PMCID: PMC6355072.

16. De la Fuente A, Mateos J, Lesende-Rodríguez I, Calamia V, Fuentes-Boquete I, de Toro FJ, et al. Proteome Analysis During Chondrocyte Differentiation in a New Chondrogenesis Model Using Human Umbilical Cord Stroma Mesenchymal Stem Cells*. Mol Cell Proteomics. 2012 Feb 1;11(2):M111.010496. doi:10.1074/mcp.M111.010496

17. Schwarzl T, Keogh A, Shaw G, Krstic A, Clayton E, Higgins DG, et al. Transcriptional profiling of early differentiation of primary human mesenchymal stem cells into chondrocytes. Sci Data. 2023 Nov 3;10(1):758. doi:10.1038/s41597-023-02686-y

18. Casorati B, Zafferri I, Castiglioni S, Maier JA. Replicative Senescence in Mesenchymal Stem Cells: An In Vitro Study on Mitochondrial Dynamics and Metabolic Alterations. Antioxidants. 2025 Apr;14(4):446. doi:10.3390/antiox14040446

19. Riss TL, Moravec RA, Duellman SJ, Niles AL. Treating Cells as Reagents to Design Reproducible Assays. SLAS Discov. 2021 Dec 1;Special Issue: Assay Guidance Manual for Drug Discovery: Robust or Go Bust26(10):1256–67. doi:10.1177/24725552211039754

20. Česnik AB, Švajger U. The issue of heterogeneity of MSC-based advanced therapy medicinal products–a review. Front Cell Dev Biol. 2024 Jul 26;12. doi:10.3389/fcell.2024.1400347

21. Altaf FM, Hering TM, Kazmi NH, Yoo JU, Johnstone B, Altaf FM, et al. Ascorbate-enhanced chondrogenesis of ATDC5 cells. Eur Cell Mater. 2006 Nov 9;12:64–70. doi:10.22203/eCM.v012a08

22. Okita K, Hikiji H, Koga A, Nagai-Yoshioka Y, Yamasaki R, Mitsugi S, et al. Ascorbic acid enhances chondrocyte differentiation of ATDC5 by accelerating insulin receptor signaling. Cell Biol Int. 2023;47(10):1737–48. doi:10.1002/cbin.12067

23. Temu TM, Wu KY, Gruppuso PA, Phornphutkul C. The mechanism of ascorbic acid-induced differentiation of ATDC5 chondrogenic cells. Am J Physiol - Endocrinol Metab. 2010 Aug;299(2):E325–34. doi:10.1152/ajpendo.00145.2010 PubMed PMID: 20530736; PubMed Central PMCID: PMC2928517.

24. Newton PT, Staines KA, Spevak L, Boskey AL, Teixeira CC, Macrae VE, et al. Chondrogenic ATDC5 cells: An optimised model for rapid and physiological matrix mineralisation. Int J Mol Med. 2012 Nov;30(5):1187–93. doi:10.3892/ijmm.2012.1114 PubMed PMID: 22941229; PubMed Central PMCID: PMC3573767.

25. Huitema LFA, Vaandrager AB, van Weeren PR, Barneveld A, Helms JB, van de Lest CHA. The nitric oxide donor sodium nitroprusside inhibits mineralization in ATDC5 cells. Calcif Tissue Int. 2006 Mar;78(3):171–7. doi:10.1007/s00223-005-1233-y PubMed PMID: 16523220.

26. Hissnauer TN, Baranowsky A, Pestka JM, Streichert T, Wiegandt K, Goepfert C, et al. Identification of molecular markers for articular cartilage. Osteoarthritis Cartilage. 2010 Dec 1;18(12):1630–8. doi:10.1016/j.joca.2010.10.002

27. Wilhelm D, Kempf H, Bianchi A, Vincourt JB. ATDC5 cells as a model of cartilage extracellular matrix neosynthesis, maturation and assembly. J Proteomics. 2020 May 15;219:103718. doi:10.1016/j.jprot.2020.103718

28. Sherman BT, Hao M, Qiu J, Jiao X, Baseler MW, Lane HC, et al. DAVID: a web server for functional enrichment analysis and functional annotation of gene lists (2021 update). Nucleic Acids Res. 2022 Jul 5;50(W1):W216–21. doi:10.1093/nar/gkac194 PubMed PMID: 35325185; PubMed Central PMCID: PMC9252805.

29. Busch T, Neubauer B, Schmitt L, Cascante I, Knoblich L, Wegehaupt O, et al. The role of the co-chaperone DNAJB11 in polycystic kidney disease: Molecular mechanisms and cellular origin of cyst formation. FASEB J. 2024;38(21):e70162. doi:10.1096/fj.202401763R

30. Demichev V, Messner CB, Vernardis SI, Lilley KS, Ralser M. DIA-NN: neural networks and interference correction enable deep proteome coverage in high throughput. Nat Methods. 2020 Jan;17(1):41–4. doi:10.1038/s41592-019-0638-x

31. Ritchie ME, Phipson B, Wu D, Hu Y, Law CW, Shi W, et al. limma powers differential expression analyses for RNA-sequencing and microarray studies. Nucleic Acids Res. 2015 Apr 20;43(7):e47. doi:10.1093/nar/gkv007

32. Heberle H, Meirelles GV, da Silva FR, Telles GP, Minghim R. InteractiVenn: a web-based tool for the analysis of sets through Venn diagrams. BMC Bioinformatics. 2015 May 22;16(1):169. doi:10.1186/s12859-015-0611-3

33. Tare RS, Howard D, Pound JC, Roach HI, Oreffo ROC. Tissue engineering strategies for cartilage generation—Micromass and three dimensional cultures using human chondrocytes and a continuous cell line. Biochem Biophys Res Commun. 2005 Jul 29;333(2):609–21. doi:10.1016/j.bbrc.2005.05.117

34. Paten JA, Martin CL, Wanis JT, Siadat SM, Figueroa-Navedo AM, Ruberti JW, et al. Molecular Interactions between Collagen and Fibronectin: A Reciprocal Relationship that Regulates *De Novo* Fibrillogenesis. Chem. 2019 Aug 8;5(8):2126–45. doi:10.1016/j.chempr.2019.05.011

35. Engvall E, Ruoslahti E. Binding of soluble form of fibroblast surface protein, fibronectin, to collagen. Int J Cancer. 1977 Jul 15;20(1):1–5. doi:10.1002/ijc.2910200102 PubMed PMID: 903179.

36. Hodax JK, Quintos JB, Gruppuso PA, Chen Q, Desai S, Jayasuriya CT. Aggrecan is required for chondrocyte differentiation in ATDC5 chondroprogenitor cells. PLOS ONE. 2019 Jun 17;14(6):e0218399. doi:10.1371/journal.pone.0218399

37. Chen L, Fink T, Zhang XY, Ebbesen P, Zachar V. Quantitative transcriptional profiling of ATDC5 mouse progenitor cells during chondrogenesis. Differentiation. 2005 Sep 1;73(7):350–63. doi:10.1111/j.1432-0436.2005.00038.x

38. Shukunami C, Ishizeki K, Atsumi T, Ohta Y, Suzuki F, Hiraki Y. Cellular Hypertrophy and Calcification of Embryonal Carcinoma-Derived Chondrogenic Cell Line ATDC5 In Vitro. J Bone Miner Res. 1997;12(8):1174–88. doi:10.1359/jbmr.1997.12.8.1174

39. Shukunami C, Ohta Y, Sakuda M, Hiraki Y. Sequential progression of the differentiation program by bone morphogenetic protein-2 in chondrogenic cell line ATDC5. Exp Cell Res. 1998 May 25;241(1):1–11. doi:10.1006/excr.1998.4045 PubMed PMID: 9633508.

40. Bartoletti G, Dong C, Umar M, He F. Pdgfra regulates multipotent cell differentiation towards chondrocytes via inhibiting Wnt9a/beta-catenin pathway during chondrocranial cartilage development. Dev Biol. 2020 Oct 1;466(1–2):36–46. doi:10.1016/j.ydbio.2020.08.004 PubMed PMID: 32800757; PubMed Central PMCID: PMC7494641.

41. Hodgkinson CP, Naidoo V, Patti KG, Gomez JA, Schmeckpeper J, Zhang Z, et al. Abi3bp is a multifunctional autocrine/paracrine factor that regulates mesenchymal stem cell biology. Stem Cells Dayt Ohio. 2013 Aug;31(8):1669–82. doi:10.1002/stem.1416 PubMed PMID: 23666637; PubMed Central PMCID: PMC3775980.

42. Sive JI, Baird P, Jeziorsk M, Watkins A, Hoyland JA, Freemont AJ. Expression of chondrocyte markers by cells of normal and degenerate intervertebral discs. Mol Pathol. 2002 Apr;55(2):91–7. doi:10.1136/mp.55.2.91 PubMed PMID: 11950957; PubMed Central PMCID: PMC1187156.

43. Lawrence JEG, Woods S, Roberts K, Sumanaweera D, Balogh P, Li T, et al. Single-cell transcriptomics identifies chondrocyte differentiation dynamics in vivo and in vitro. Dev Cell. 2025 Nov 17;60(22):3066–3084.e8. doi:10.1016/j.devcel.2025.06.031

44. Sebastian A, McCool JL, Hum NR, Murugesh DK, Wilson SP, Christiansen BA, et al. Single-Cell RNA-Seq Reveals Transcriptomic Heterogeneity and Post-Traumatic Osteoarthritis-Associated Early Molecular Changes in Mouse Articular Chondrocytes. Cells. 2021 Jun;10(6):1462. doi:10.3390/cells10061462

45. Zhao Q, Eberspaecher H, Lefebvre V, De Crombrugghe B. Parallel expression of Sox9 and Col2a1 in cells undergoing chondrogenesis. Dev Dyn Off Publ Am Assoc Anat. 1997 Aug;209(4):377–86. doi:10.1002/(SICI)1097-0177(199708)209:4<377::AID-AJA5>3.0.CO;2-F PubMed PMID: 9264261.

46. Gu J, Lu Y, Li F, Qiao L, Wang Q, Li N, et al. Identification and characterization of the novel Col10a1 regulatory mechanism during chondrocyte hypertrophic differentiation. Cell Death Dis. 2014 Oct;5(10):e1469. doi:10.1038/cddis.2014.444 PubMed PMID: 25321476; PubMed Central PMCID: PMC4649528.

47. Holm E, Gleberzon JS, Liao Y, Sørensen ES, Beier F, Hunter GK, et al. Osteopontin mediates mineralization and not osteogenic cell development in vitro. Biochem J. 2014 Dec 5;464(3):355–64. doi:10.1042/BJ20140702

48. Surmann-Schmitt C, Dietz U, Kireva T, Adam N, Park J, Tagariello A, et al. Ucma, a novel secreted cartilage-specific protein with implications in osteogenesis. J Biol Chem. 2008 Mar 14;283(11):7082–93. doi:10.1074/jbc.M702792200 PubMed PMID: 18156182.

49. Akiyama H, Chaboissier MC, Martin JF, Schedl A, de Crombrugghe B. The transcription factor Sox9 has essential roles in successive steps of the chondrocyte differentiation pathway and is required for expression of Sox5 and Sox6. Genes Dev. 2002 Nov 1;16(21):2813–28. doi:10.1101/gad.1017802 PubMed PMID: 12414734; PubMed Central PMCID: PMC187468.

50. Lefebvre V, Dvir-Ginzberg M. SOX9 and the many facets of its regulation in the chondrocyte lineage. Connect Tissue Res. 2017 Jan;58(1):2–14. doi:10.1080/03008207.2016.1183667 PubMed PMID: 27128146; PubMed Central PMCID: PMC5287363.

51. Ziros PG, Basdra EK, Papavassiliou AG. Runx2: of bone and stretch. Int J Biochem Cell Biol. 2008;40(9):1659–63. doi:10.1016/j.biocel.2007.05.024 PubMed PMID: 17656144.

52. Caron MMJ, Eveque M, Cillero-Pastor B, Heeren RMA, Housmans B, Derks K, et al. Sox9 Determines Translational Capacity During Early Chondrogenic Differentiation of ATDC5 Cells by Regulating Expression of Ribosome Biogenesis Factors and Ribosomal Proteins. Front Cell Dev Biol. 2021 Jun 21;9:686096. doi:10.3389/fcell.2021.686096 PubMed PMID: 34235151; PubMed Central PMCID: PMC8256280.

53. Dong X, Xu X, Yang C, Luo Y, Wu Y, Wang J. USP7 regulates the proliferation and differentiation of ATDC5 cells through the Sox9-PTHrP-PTH1R axis. Bone. 2021 Feb 1;143:115714. doi:10.1016/j.bone.2020.115714

54. Leung VYL, Gao B, Leung KKH, Melhado IG, Wynn SL, Au TYK, et al. SOX9 Governs Differentiation Stage-Specific Gene Expression in Growth Plate Chondrocytes via Direct Concomitant Transactivation and Repression. PLOS Genet. 2011 Nov 3;7(11):e1002356. doi:10.1371/journal.pgen.1002356

55. Kim IS, Otto F, Zabel B, Mundlos S. Regulation of chondrocyte differentiation by Cbfa1. Mech Dev. 1999 Feb;80(2):159–70. doi:10.1016/s0925-4773(98)00210-x PubMed PMID: 10072783.

56. Horkay F, Basser PJ, Hecht AM, Geissler E. Structure and Properties of Cartilage Proteoglycans. Macromol Symp. 2017;372(1):43–50. doi:10.1002/masy.201700014

57. Kiani C, Chen L, Wu YJ, Yee AJ, Yang BB. Structure and function of aggrecan. Cell Res. 2002 Mar;12(1):19–32. doi:10.1038/sj.cr.7290106

58. Neame PJ, Barry FP. The link proteins. Experientia. 1993 May 1;49(5):393–402. doi:10.1007/BF01923584

59. Vidak E, Javoršek U, Vizovišek M, Turk B. Cysteine Cathepsins and Their Extracellular Roles: Shaping the Microenvironment. Cells. 2019 Mar 20;8(3):264. doi:10.3390/cells8030264 PubMed PMID: 30897858; PubMed Central PMCID: PMC6468544.

60. Shao X, Gomez CD, Kapoor N, Considine JM, Grams C, Gao Y (Tom), et al. MatrisomeDB 2.0: 2023 updates to the ECM-protein knowledge database. Nucleic Acids Res. 2023 Jan 6;51(D1):D1519–30. doi:10.1093/nar/gkac1009

61. Alcaide-Ruggiero L, Molina-Hernández V, Granados MM, Domínguez JM. Main and Minor Types of Collagens in the Articular Cartilage: The Role of Collagens in Repair Tissue Evaluation in Chondral Defects. Int J Mol Sci. 2021 Dec 11;22(24):13329. doi:10.3390/ijms222413329 PubMed PMID: 34948124; PubMed Central PMCID: PMC8706311.

62. Georgieva VS, Etich J, Bluhm B, Zhu M, Frie C, Wilson R, et al. Ablation of the miRNA Cluster 24 Has Profound Effects on Extracellular Matrix Protein Abundance in Cartilage. Int J Mol Sci. 2020 Jan;21(11):4112. doi:10.3390/ijms21114112

63. Bubb K, Holzer T, Nolte JL, Krüger M, Wilson R, Schlötzer-Schrehardt U, et al. Mitochondrial respiratory chain function promotes extracellular matrix integrity in cartilage. J Biol Chem. 2021 Sep 22;297(4):101224. doi:10.1016/j.jbc.2021.101224 PubMed PMID: 34560099; PubMed Central PMCID: PMC8503590.

64. Chen L, Fink T, Ebbesen P, Zachar V. Temporal transcriptome of mouse ATDC5 chondroprogenitors differentiating under hypoxic conditions. Exp Cell Res. 2006 Jun 10;312(10):1727–44. doi:10.1016/j.yexcr.2006.02.013 PubMed PMID: 16580664.

65. Yuan Z, Liu S, Song W, Liu Y, Bi G, Xie R, et al. Galactose Enhances Chondrogenic Differentiation of ATDC5 and Cartilage Matrix Formation by Chondrocytes. Front Mol Biosci. 2022 May 9;9. doi:10.3389/fmolb.2022.850778

66. Farcasanu MV, Ruiz T de las H, Brito FMJ de S, Soul J, Coxhead J, German MJ, et al. Dynamic Compression Improves Chondrogenesis in the Tissue Engineered Model of Cartilage. Biotechnol Bioeng. 2025;122(9):2574–91. doi:10.1002/bit.29026

67. Challa TD, Rais Y, Ornan EM. Effect of adiponectin on ATDC5 proliferation, differentiation and signaling pathways. Mol Cell Endocrinol. 2010 Jul 29;323(2):282–91. doi:10.1016/j.mce.2010.03.025

68. ATDC5 (ECACC 99072806) Growth Profile, accessed 28.01.2026 [Internet]. 2026 [cited 2026 Jan 28]. Available from: https://www.culturecollections.org.uk/nop/product/atdc5

69. Gottardi R, Hansen U, Raiteri R, Loparic M, Düggelin M, Mathys D, et al. Supramolecular Organization of Collagen Fibrils in Healthy and Osteoarthritic Human Knee and Hip Joint Cartilage. PLOS ONE. 2016 Oct 25;11(10):e0163552. doi:10.1371/journal.pone.0163552

70. Moger CJ, Barrett R, Bleuet P, Bradley DA, Ellis RE, Green EM, et al. Regional variations of collagen orientation in normal and diseased articular cartilage and subchondral bone determined using small angle X-ray scattering (SAXS). Osteoarthritis Cartilage. 2007 Jun 1;15(6):682–7. doi:10.1016/j.joca.2006.12.006

71. Molin AN, Contentin R, Angelozzi M, Karvande A, Kc R, Haseeb A, et al. Skeletal growth is enhanced by a shared role for SOX8 and SOX9 in promoting reserve chondrocyte commitment to columnar proliferation. Proc Natl Acad Sci. 2024 Feb 20;121(8):e2316969121. doi:10.1073/pnas.2316969121

72. Wang Y, Middleton F, Horton JA, Reichel L, Farnum CE, Damron TA. Microarray analysis of proliferative and hypertrophic growth plate zones identifies differentiation markers and signal pathways. Bone. 2004 Dec 1;35(6):1273–93. doi:10.1016/j.bone.2004.09.009

73. Tchetina E, Mwale F, Poole A. Distinct Phases of Coordinated Early and Late Gene Expression in Growth Plate Chondrocytes in Relationship to Cell Proliferation, Matrix Assembly, Remodeling, and Cell Differentiation. J Bone Miner Res. 2003;18(5):844–51. doi:10.1359/jbmr.2003.18.5.844

74. Makhijani P, Basso PJ, Chan YT, Chen N, Baechle J, Khan S, et al. Regulation of the immune system by the insulin receptor in health and disease. Front Endocrinol. 2023 Mar 13;14:1128622. doi:10.3389/fendo.2023.1128622 PubMed PMID: 36992811; PubMed Central PMCID: PMC10040865.

75. Luo Z, Tian M, Yang G, Tan Q, Chen Y, Li G, et al. Hypoxia signaling in human health and diseases: implications and prospects for therapeutics. Signal Transduct Target Ther. 2022 Jul 7;7(1):218. doi:10.1038/s41392-022-01080-1

76. Cummins EP, Berra E, Comerford KM, Ginouves A, Fitzgerald KT, Seeballuck F, et al. Prolyl hydroxylase-1 negatively regulates IkappaB kinase-beta, giving insight into hypoxia-induced NFkappaB activity. Proc Natl Acad Sci U S A. 2006 Nov 28;103(48):18154–9. doi:10.1073/pnas.0602235103 PubMed PMID: 17114296; PubMed Central PMCID: PMC1643842.

77. Liang T, Li P, Liang A, Zhu Y, Qiu X, Qiu J, et al. Identifying the key genes regulating mesenchymal stem cells chondrogenic differentiation: an in vitro study. BMC Musculoskelet Disord. 2022 Nov 15;23(1):985. doi:10.1186/s12891-022-05958-7

78. Zhou J, Li C, Yu A, Jie S, Du X, Liu T, et al. Bioinformatics analysis of differentially expressed genes involved in human developmental chondrogenesis. Medicine (Baltimore). 2019 Jul;98(27):e16240. doi:10.1097/MD.0000000000016240

79. Rankin EB, Giaccia AJ, Schipani E. A Central Role for Hypoxic Signaling in Cartilage, Bone, and Hematopoiesis. Curr Osteoporos Rep. 2011 Jun;9(2):46–52. doi:10.1007/s11914-011-0047-2 PubMed PMID: 21360287; PubMed Central PMCID: PMC4012534.

80. Wagener R, Ehlen HWA, Ko YP, Kobbe B, Mann HH, Sengle G, et al. The matrilins – adaptor proteins in the extracellular matrix. FEBS Lett. 2005 Jun 13;Budapest Special Issue579(15):3323–9. doi:10.1016/j.febslet.2005.03.018

81. Posey KL, Coustry F, Hecht JT. Cartilage oligomeric matrix protein: COMPopathies and beyond. Matrix Biol J Int Soc Matrix Biol. 2018 Oct;71–72:161–73. doi:10.1016/j.matbio.2018.02.023 PubMed PMID: 29530484; PubMed Central PMCID: PMC6129439.

82. Yang CY, Chanalaris A, Troeberg L. ADAMTS and ADAM metalloproteinases in osteoarthritis – looking beyond the ‘usual suspects.’ Osteoarthritis Cartilage. 2017 Jul;25(7):1000–9. doi:10.1016/j.joca.2017.02.791 PubMed PMID: 28216310; PubMed Central PMCID: PMC5473942.

