## Supplemental material for "Deep phenotyping of ATDC5-derived in vitro cartilage organoids"

Supplementary Figures and Supplementary Files legends

### Supplementary Figures

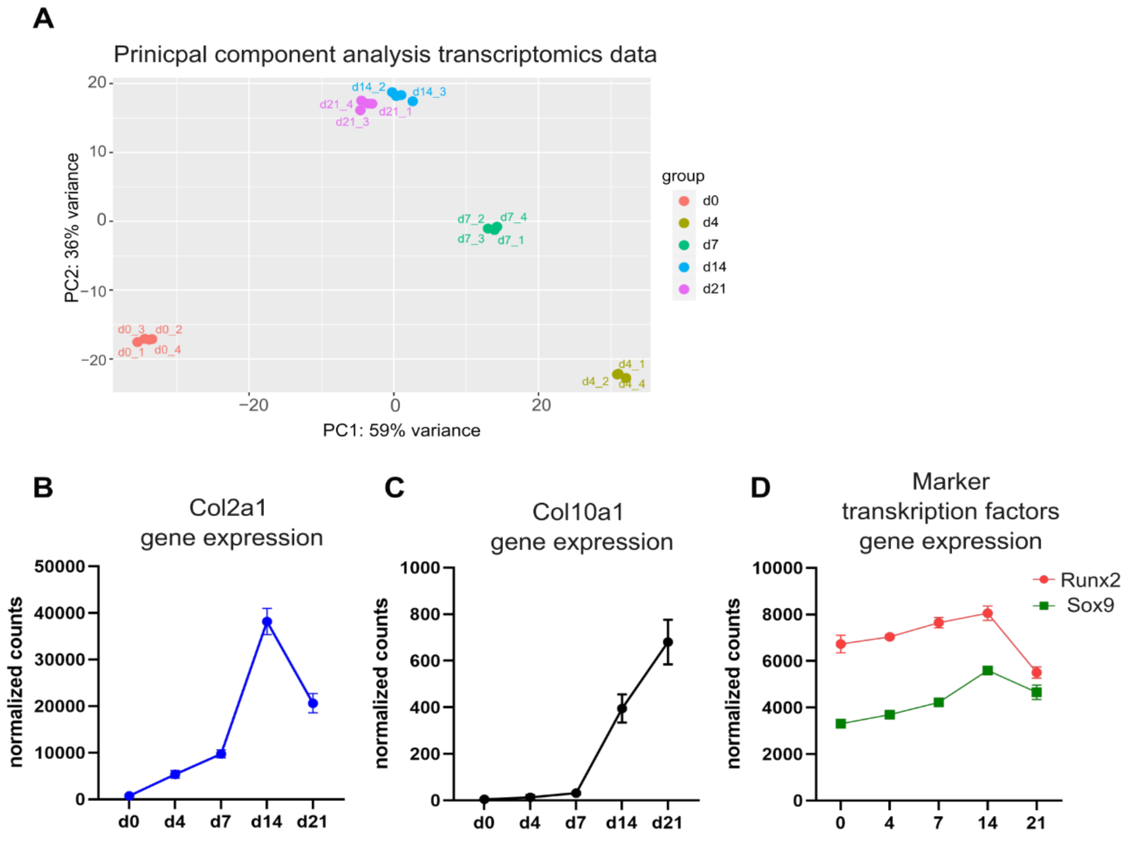


**Figure S1:**(A) Principal component analysis of transcriptomics samples based on DEseq2 normalized data shows distinct clustering of samples per time point. Samples from day 14 and day 21 cluster closely together. One sample at day 4 could not be processed and was excluded from downstream analysis; therefore, three biological replicates were analyzed for this time point and four for all others. PC1 and PC2 explain 59% and 36% of total variance. Samples are color-coded according to time point. (B-D) Line plots illustrating expression of Co2la1 (B), Col10a1 (C) and Sox9 and Runx2 (D) gene expression values derived from DESeq2-normalized RNA-seq counts. Data are presented as mean ± 3 (n=3-4).

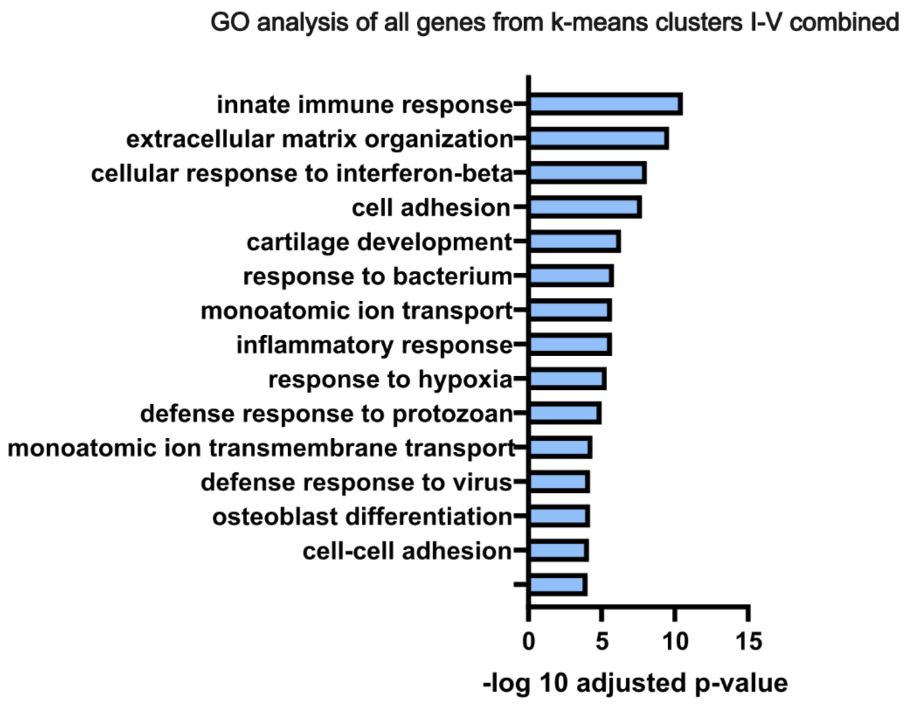


**Figure S2:** Significant GO terms "biological process" after GO analysis of all genes in clusters I-V revealed that genes that change most in expression during the differentiation process are associated with ECM and immunity linked GO terms. DAVID Gene Ontology (https://davidbioinformatics.nih.gov/) Functional Annotation Tool was used to investigate GO terms in the category "biological process".

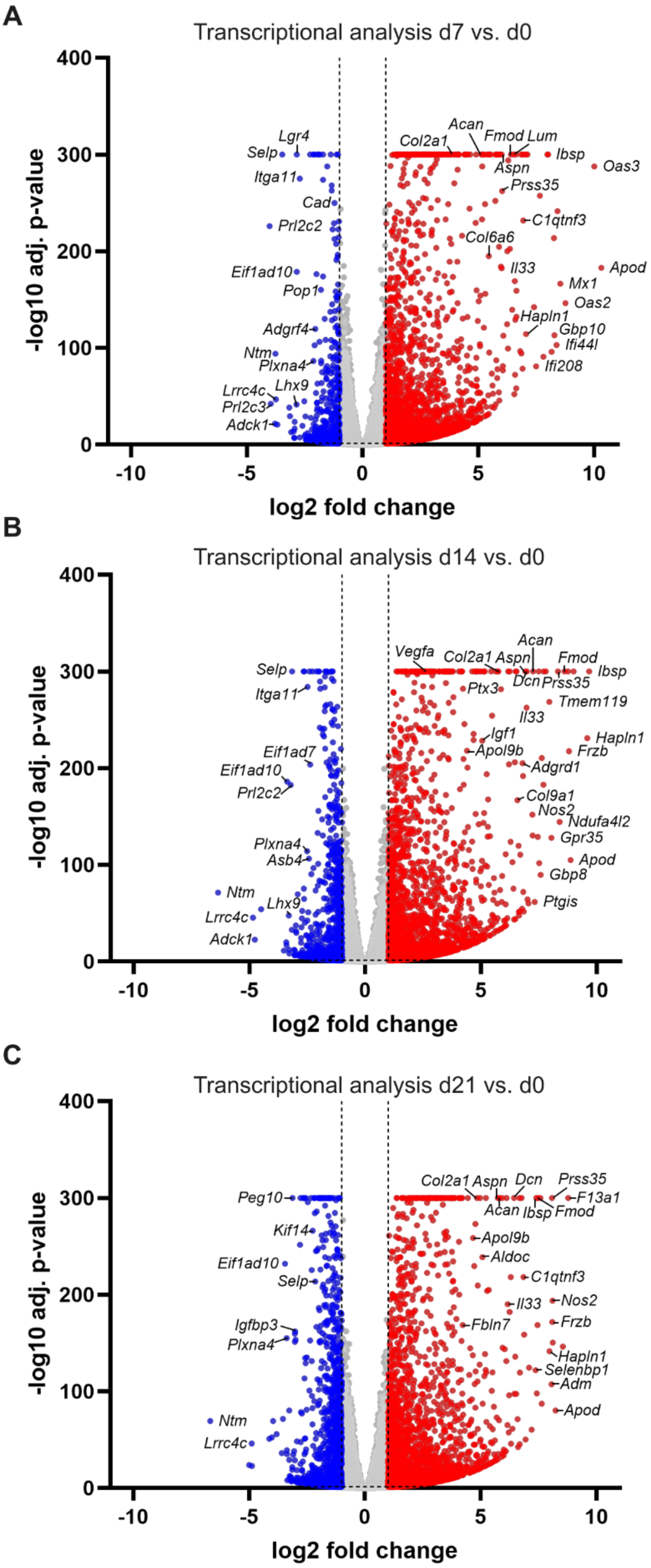


**Figure S3:** (A-C) Volcano plots illustrating transcriptional analysis of transcriptomics samples at day 7 vs. day 0 (A), day 14 vs. day 0 (B) and day 21 vs. day 0 (C), highlighting significantly increased expression of chondrocyte markers such as *Col2a1*, *Acan* and *Fmod* at all time points compared to day 0. For visualization purposes, −log10 adjusted p-values were capped at 300. Genes were color-coded according to log2fold change thresholds of ±1 and adjusted p-value ≤ 0.05.

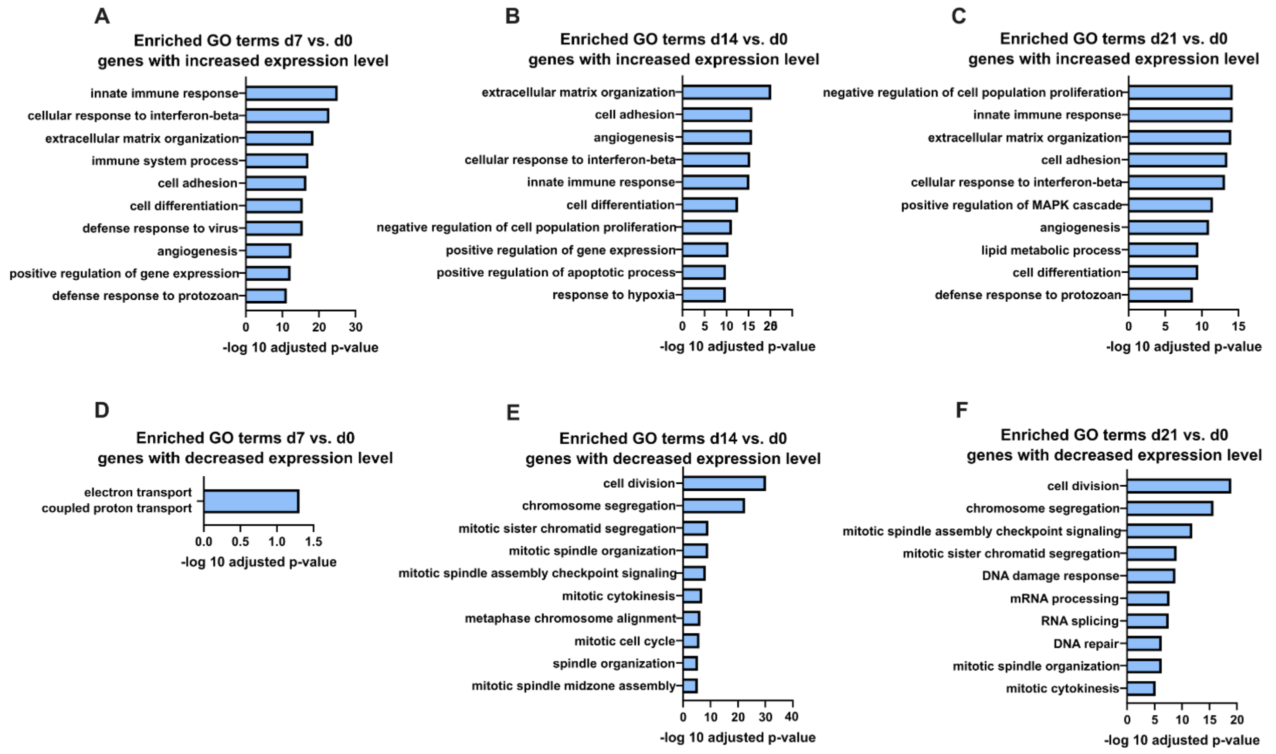


**Figure S4:**

(A-F) Significant GO terms “biological process” after GO analysis of genes with significantly (adjusted p-value ≤ 0.05, |log2 fold change| ≥ 1) increased (A-C) or decreased (D-F) expression at time points as indicated. Top 10 or all significant identified terms are displayed. For analysis of genes with decreased expression at day 7 vs. day 0 (D) and day 14 vs. day 0 (E), predicted genes were excluded.

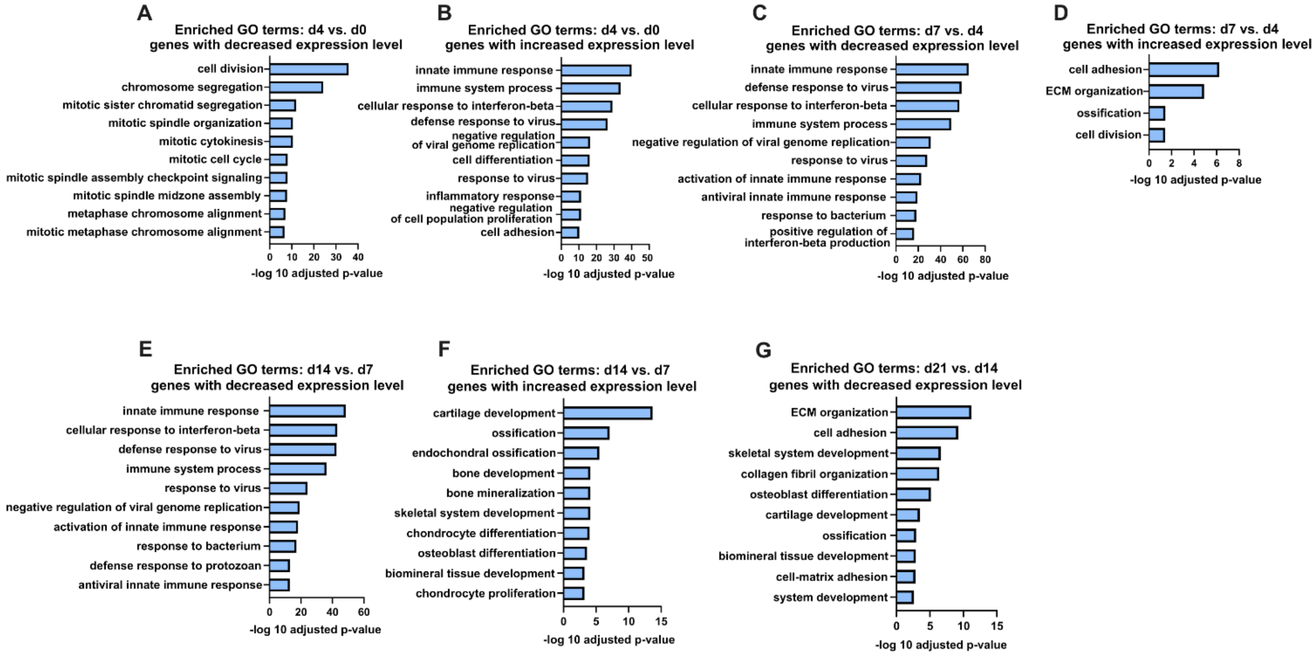


**Figure S5:**

(A-G) Significant GO terms “biological process” after GO analysis of genes with significantly (adjusted p-value ≤ 0.05, |log2 fold change| ≥ 1) decreased (A, C, E, G) or increased (B, D, F) expression at consecutive time points as indicated. Top 10 or all significant terms are displayed. For analysis of genes with decreased expression at day 4 vs. day 0, predicted genes were excluded. The analysis of genes with increased expression at day 21 vs. day 14 did not reveal any significant GO terms (Supplementary File S3).

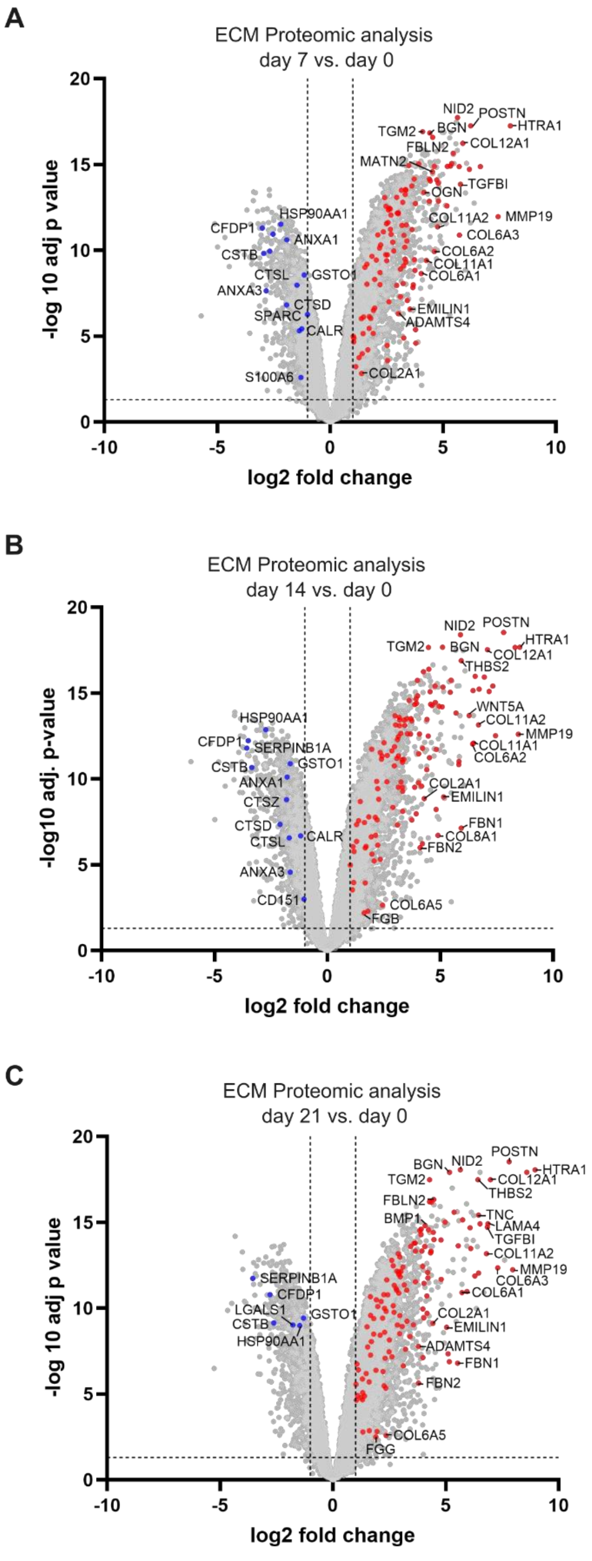


**Figure S6:**

Volcano plots illustrating proteomic analysis of samples at day 7 vs. day 0 (A), day 14 vs. day 0 (B) and day 21 vs. day 0 (C) reveals increased expression of ECM proteins annotated in

GO:0031012 ECM over time. Proteins not annotated in GO:0031012 ECM are displayed in grey. ECM proteins were color-coded according to adjusted $p \leq 0.05$ and log2fold change thresholds of $\pm 1$.

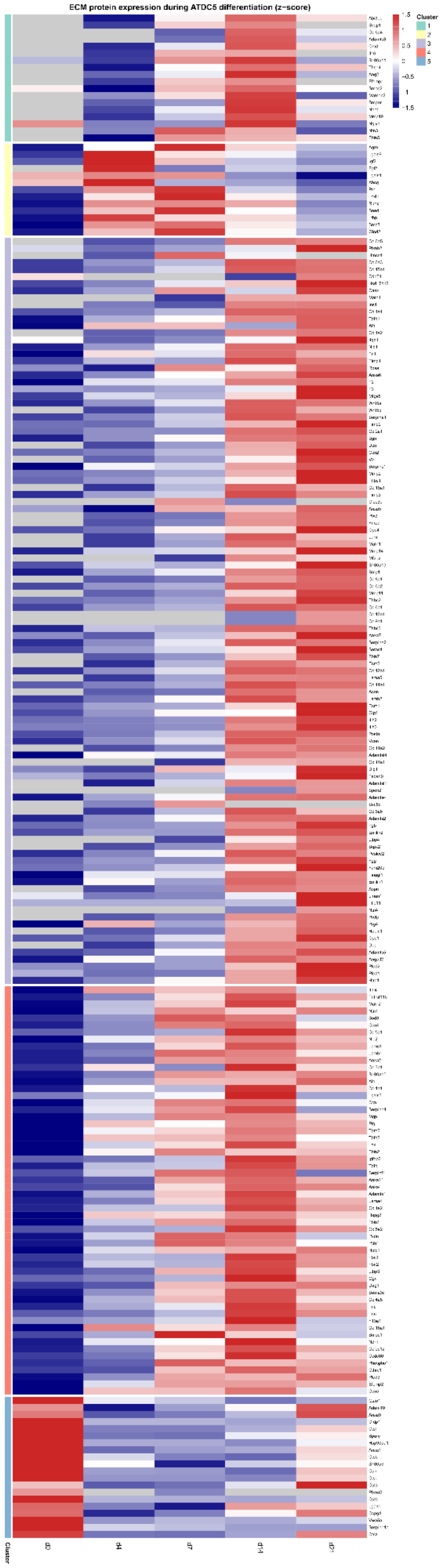


**Figure S7:** K-means clustered heatmap of 216 identified ECM proteins annotated in GO:0031012 Extracellular matrix (k=5). The majority of proteins show increasing expression over the course of the differentiation, except for proteins in cluster 2 and 5. Normalized expression values derived from the limma input dataset were used to calculate z-scores. Expression values are scaled per protein (z-score) to illustrate relative abundance patterns across timepoints.

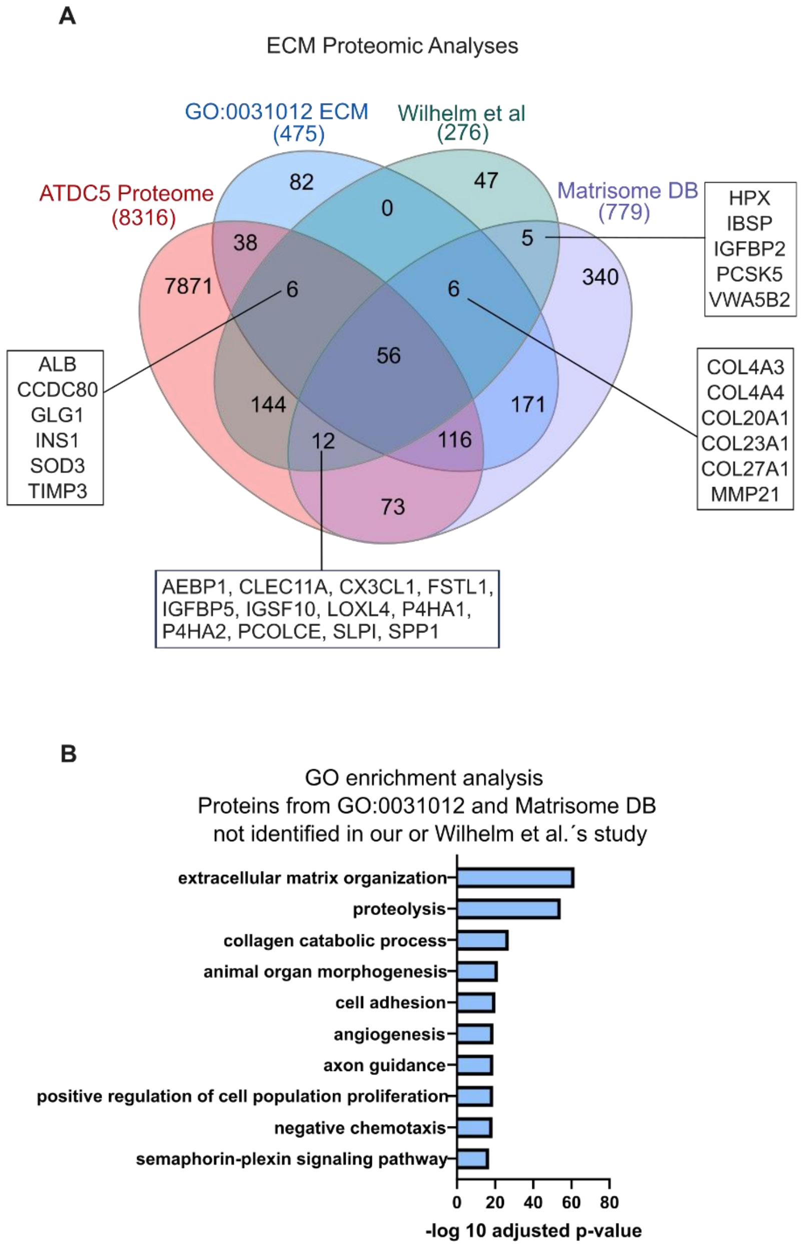


**Figure S8:**

(A) Venn Diagram illustrating overlap of proteins identified in our study, Wilhelm et al´s study, GO:0031012 (extracellular matrix) and murine cartilage proteins from the matrisome database. (B) GO analysis of “biological process” with proteins listed in GO:0031012 (extracellular matrix) and/or murine cartilage proteins from the matrisome database, but not identified in neither our nor Wilhelm et al.´s study (n=593), indicating that a large subset of not identified proteins are associated with proteolytic and collagen catabolic functions.”

**Captions for Supplementary Files**

**Supplementary File S1**: Names of gene clusters, genes sorted for clusters and GO analysis for each cluster

**Supplementary File S2:** DEseq2 result files for transcriptomic analyses presented

**Supplementary File S3:** Full list of GO enrichment analysis “biological process” results for all GO analyses presented, separated for genes with decreasing or increasing expression

**Supplementary File S4:** Table with proteins identified in our study at min. 2 time points at min one time point with normalized expression value

**Supplementary File S5:** Lists of intersections from Venn diagrams displayed in Figures 6 and S7

**Supplementary File S6:** Full list of GO enrichment analysis “biological process” with proteins listed in GO:0031012 (extracellular matrix) and/or murine cartilage proteins from the matrisome database, but not identified in neither our nor Wilhelm et al.´s study, displayed in Figure S7.
